# Characterisation and phylogenetic analysis of Murray Valley encephalitis virus (MVEV) from the Ralph Doherty collection

**DOI:** 10.1101/2025.07.24.666511

**Authors:** Freyja E. Stam, Cameron R. Bishop, Bing Tang, Abigail L. Cox, Kexin Yan, Andrii Slonchak, Jessica J. Harrison, Jody Hobson-Peters, Roy A. Hall, Gorben P. Pijlman, Daniel J. Rawle, Andreas Suhrbier, Wilson Nguyen

## Abstract

Murray Valley encephalitis virus (MVEV) is a zoonotic flavivirus endemic to Australia and Papua New Guinea. A recent outbreak of MVEV has prompted renewed concerns regarding the potential for MVEV to generate disease outbreaks. Currently, nine full length sequences of MVEV are publicly available, divided into four genotypes (G1-G4). Herein, we sequenced MVEV isolates from the Ralph Doherty Virus Collection, a virus bank with Australian field isolates dating between the 1950s-1980s, and determined their phylogenetic relationship with existing isolates to provide insights into virus evolution and genetic diversity. Additionally, we characterised isolates from different genotypes both *in vitro* using human neuronal cells, and *in vivo* using C57BL/6J mice, to provide additional insight into MVEV pathogenicity and establish models of MVEV disease that recapitulate MVEV human disease. We found 15 new full length sequences of MVEV, which primarily clustered into the dominant genotype, G1. Additionally, we show MVEV can be lethal and neuroinvasive in C57BL/6J mice, recapitulating histological lesions identified in human infection. Overall, our study contributes significant genomic sequences to the current MVEV database and establishes mouse models of disease and infection which can be used for mechanistic studies and evaluation of new interventions.

## Introduction

Murray Valley encephalitis virus (MVEV) is a single-stranded positive-sense RNA virus from the *Flaviviridae* family belonging to the Japanese encephalitis serocomplex, alongside other encephalitic flaviviruses like Japanese encephalitis virus (JEV) and West Nile virus (WNV). MVEV exists in a zoonotic cycle between waterbirds and *Culex* mosquitoes, which can transmit the virus to humans. Most human cases of MVEV remain asymptomatic, yet an estimated 1:150 to 1:1000 cases develop clinical encephalitis, with symptoms ranging from fever and headache to neurological deterioration (Knox et al., 2012). Of these symptomatic cases, 15-30% are fatal and another 30-50% retains neurological symptoms (Selvey et al., 2014a). Aside from humans, MVEV can also cause fatal encephalitis in horses (Gordon et al., 2012; Roche et al., 2013).

MVEV was first isolated in 1951 in the Murray Valley region and has remained endemic to Australia and Papua New-Guinea. Lately, the virus has become increasingly active in more populous areas of Australia. In 2011, widespread activity of the virus in humans was seen for the first time since the nationwide epidemic in 1974, resulting in seventeen human cases (Selvey et al., 2014b). The most recent outbreak occurred in the summer of 2022-2023 reported 27 clinical cases across Australia (National Notifiable Diseases Surveillance System, 2025). While these outbreaks raise concerns over the increasing spread of MVEV, no licensed treatment or vaccines are available. Efforts to develop such treatments for MVEV neuropathology are challenging due to the difficulties in diagnosing early infection before the virus has infected the brain, and once the brain is infected, such therapeutics need to effectively enter the brain. While a range of MVEV mouse models have been reported (Supplementary Table 1), many of these studies focus solely on mortality post infection without examining patterns that recapitulate MVEV disease in humans. In addition, these studies focus on the prototype strain of MVEV, 1-51, which belongs to genotype 1.

Four genotypes (G1-G4) of MVEV have previously been identified, based on genetic analyses including RNase oligonucleotide mapping and genome sequencing of the pre-membrane and envelope region (Coelen and Mackenzie, 1988, 1990; Johansen et al., 2007; Lobigs et al., 1988; Poidinger et al., 1996; Williams et al., 2015). These studies showed G1 as the dominant genotype, with a majority of existing MVEV isolates clustering in this clade, including strains from recent MVEV outbreaks. While some phylogenetic analyses on MVEV sequences have been performed, these studies have focused on partial genome sequences within the pre-membrane, envelope, and 5’ UTR region or on RNase oligonucleotide mapping (Coelen and Mackenzie, 1988, 1990; Johansen et al., 2007; Lobigs et al., 1988; Poidinger et al., 1996; Williams et al., 2015). To the best of our knowledge, only 9 unique full length genome sequences of MVEV are publicly available, creating challenges in understanding evolutionary changes of the virus which may influence its pathogenicity and virulence. In addition, characterisation of other genotypes (other than G1) both *in vitro* and *in vivo* is limited (Supplementary Table 1).

In this study, we performed an expanded phylogenetic analysis on the full-length genome sequences of MVEV, including 15 new unique isolates from the Ralph Doherty virus collection (an Australian virus bank that contains field isolates collected between the 1950s to the 1980s), to further understand the genetic relationships between MVEV strains. Additionally, *in vitro* and *in vivo* characterisation of different MVEV isolates was performed to establish models of infection and disease for mechanistic studies.

## Materials and Methods

### Ethics statement

All mouse work was conducted in accordance with the “Australian code for the care and use of animals for scientific purposes” as defined by the National Health and Medical Research Council of Australia. Mouse work was approved by the QIMR Berghofer Medical Research Institute (MRI) Animal Ethics Committee (P3746, A2109-612).

### Cell lines and culture

Vero E6 (C1008, ECACC, Wiltshire, England; obtained via Sigma-Aldrich, St. Louis, MO, USA), BHK-21 (ATCC# CCL-10) and C6/36 cells (ATCC# CRL-1660) were cultured in Roswell Park Memorial Institute (RPMI) 1640 medium, supplemented with 10% fetal bovine serum (FBS), penicillin (100 IU/ml)/streptomycin (100 µg/ml) (Gibco/Life Technologies) and L-glutamine (2 mM) (Life Technologies). PK-15 (ATCC# CCL-33) cells were cultured in Eagle’s Minimum Essential Medium (EMEM), supplemented with 10% FBS and penicillin (100 IU/ml)/streptomycin (100 µg/ml) (Gibco/Life Technologies). Vero E6, BHK-21, and PK-15 cells were cultured at 37 °C and 5% CO_2_, and C6/36 cells were cultured at 27 °C and 5% CO_2_.

RENcell VM Human Neural Progenitor Cell Line (Sigma-Aldrich) were cultured in medium comprising DMEM F-12 (Thermo Fisher Scientific), penicillin (100 IU/ml)/streptomycin (100 μg/ml) (Gibco/Life Technologies), 20 ng/ml FGF (STEMCELL Technologies), 20 ng/ml EGF (STEMCELL Technologies), and 2% B27 supplements (Thermo Fisher Scientific). Cells were detached using StemPro Accutase Cell Dissociation Reagent (Thermo Fisher Scientific), and were cultured in Matrigel (Bio-Strategy). To produce mature neurons, cells were seeded into poly-L-ornithine-(Merck) and laminin-coated (Thermo Fisher Scientific) 12-wells plate at 3.5 × 10^4^/ml. After incubation of 48 hours at 37 °C, Stemcell media was replaced with Neural Differentiation media comprising Neurobasal media (Thermo Fisher Scientific), 2% B27 (Thermo Fisher Scientific), 1% GlutaMAX (Thermo Fisher Scientific), and 1% insulin-transferrin-selenium (Thermo Fisher Scientific). Neurobasal media was replaced every 2-4 days after washing with Dulbecco’s Phosphate-Buffered Saline (DPBS) (Thermo Fisher Scientific).

For crystal violet staining of remaining cells after infection, formaldehyde (7.5% w/v)/crystal violet (0.05% w/v) was added to wells overnight, plates washed twice in water, and plates dried overnight.

### Virus isolates and culture

The Ralph Doherty Virus collection located at QIMR Berghofer Medical Research Institute houses a significant number of virus isolates from field studies dated between the 1950s-1980s. The collection contained 48 unique virus isolates (Supplementary Table 2), of which 43 caused cytopathic effects (CPE) in BHK cells and were sequenced. MVEV CY1189, MVEV 6684 and MVEV OR156 (which was originally in the Doherty collection but unable to be revived) were generously donated by Prof Roy Hall and Dr Jody-Hobson Peters from the University of Queensland.

Virus stocks were generated by infection of either C6/36 cells or PK-15 cells at MOI ≈ 0.1, with supernatant collected after ∼5 days, cell debris removed by centrifugation at 3000 x *g* for 15 min at 4 °C, and virus aliquoted and stored at – 80 °C. Virus stocks used in these experiments underwent less than three passages in our laboratory. Data on prior available passage history of these isolates is unavailable. Virus titres were determined using standard CCID_50_ assays (see below); results can be found in Supplementary Table 3.

### Validation of virus stock sequences

Viral RNA (MVEV CY1189, MVEV 6684 and MVEV OR156) was extracted from infected cells using TRIzol (Invitrogen) as per manufacturer’s instructions. cDNA was synthesized using ProtoScript First Strand cDNA Synthesis Kit (New England Biolabs) as per manufacturer’s instructions. PCR was performed using Q5 High-Fidelity 2X Master Mix (New England Biolabs) as per manufacturer’s instructions with the following primers; MVEV OR156 PrM Forward 5’ GTAAGATAATGATGACCGTGAACGCCACGGAC 3’ and Envelope Reverse 5’ CGGCAAGAGCAAGTCATTAAACCATTCGCG 3’. PCR products were run on a 1% agarose gel to confirm results. RT-qPCR was also performed using iTaq Universal One-Step RT-qPCR Kit (Bio-Rad) as per manufacturer’s instructions with the following primers; MVEV 1-51 PrM Forward 5’ AAGCTTTCCACCTTCCAGGGCAAGATAATGATG 3’ and Envelope Reverse 5’ TTGCAGGTGATGTCCATGGCAAGAGCAA 3’, MVEV OR156 PrM Forward 5’ GTAAGATAATGATGACCGTGAACGCCACGGAC 3’ and Envelope Reverse 5’ CGGCAAGAGCAAGTCATTAAACCATTCGCG 3’, and MVEV MK6684 PrM Forward 5’ TCCACCTTCCAGGGCAAAATAATGATGACC 3’ and Envelope Reverse 5’ GATGTCCACGGCAGGAGCAGATCATTAAACC 3’ or MVEV NG156 Envelope Reverse 5’ CTTGAAGGTGATGTCCACGGCAAGAGCAAATC 3’.

### RNA-Sequencing

Viral RNA was extracted as described above, and RNA-Seq was undertaken as described using Illumina Nextseq 550 platform generating 75 bp paired end reads (Bishop et al., 2022; Rawle et al., 2020) The per base sequence quality for >90% bases was above Q30 for all samples. Virus reads were aligned to an *Aedes albopictus* reference genome using STAR aligner. Sequences were assembled and analysed using the software, Geneious version X (Geneious, Auckland, New Zealand) using the prototype isolate MVEV 1-51 (GenBank: AF161266.1) as a reference.

### Phylogenetic analyses and tree construction

Full length or partial genome viral sequences were aligned with all publicly available sequences using the ClustalW plugin with default parameters in MEGA11 (Molecular Evolutionary Genetics Analysis 11). Maximum likelihood trees were composed with 1000 bootstrap replicates from full genome, PrME, NS1-NS5, 5’ UTR, and 3’UTR nucleotide and/or amino acid sequences using MEGA11. Trees were rooted to the phylogenetically related Alfuy MRM3929 (GenBank: AY898809) as an out-group.

### Cell culture infectious dose (CCID_50_) assays

CCID_50_ assays were performed as previously described (Nguyen et al., 2024; Nguyen et al., 2020; Rawle et al., 2020). C6/36 cells were plated in 96 well flat bottom plates at 2 × 10^5^ cells per well in 100 µl of medium. For tissue titrations, tissues were homogenized in tubes each containing 4 ceramic beads twice at 6000 × *g* for 30 s, followed by centrifugation twice at 21,000 × *g* for 5 min. Samples underwent 10-fold serial dilutions in 100 µl RPMI 1640 supplemented with 2% FBS, performed in quadruplicate for tissue homogenate and in duplicate for serum. For mouse tissues and serum, a volume of 100 µl of serially diluted samples was added to each well of 96 well plates containing C6/36 cells, and the plates were cultured for 5 days at 37 °C and 5% CO_2_. 25 µl of supernatant from infected C6/36 cells were then passaged on to Vero E6 cells plated the day before at 2 × 10^5^ cells per well in 96 well flat bottom plates. Supernatants from in vitro stock cultures were both titrated directly onto Vero E6 cells and passaged from C6/36 onto Vero E6 cells as described. Vero E6 cells were cultured for 5 days cytopathic effect was scored, and the virus titre was calculated using the method of Spearman and Karber (Ramakrishnan, 2016)

### Mouse infections

Female C57BL/6J mice were purchased from Australian BioResources (Moss Vale, NSW, Australia). Mice housing conditions were described previously (Dumenil et al., 2023). At 6 weeks old, mice were infected subcutaneously (s.c.) at the base of the tail with 100 µl of virus inoculum at 5 × 10^3^. Serum was collected via tail nicks into Microvette Serum-Gel 500 µl blood collection tubes (Sarstedt, Numbrecht, Germany). Mice were weighed and monitored for disease manifestations and were euthanized using CO_2_ based on a scorecard system (Nguyen et al., 2024). At necropsy, brain and spleens were collected for virus titrations by CCID_50_ assays and/or for histology, and blood was collected into Micro Sample tube Serum Gel 1.1 ml blood collection tubes (Sarstedt, Numbrecht, Germany) via cardiac puncture.

### Histopathology and immunohistochemistry

Brains were fixed in 10% formalin and embedded in paraffin. Sections were stained using H&E (Sigma-Aldrich) and slides were scanned using Aperio AT Turbo (Aperio, Vista, CA USA) and analysed using Aperio ImageScope software v10 (LeicaBiosystems, Mt Waverley, Australia). Leukocyte infiltrates were quantified by measuring nuclear (strong purple staining)/cytoplasmic (total red staining) pixel rations in scanned H&E stained images and undertaken using Aperio Positive Pixel Count Algorithm (Leica Biosystems). Sections were stained for anti-flavivirus non-structural protein 1 (NS1) immunohistochemistry using 4G4 as previously described (Nguyen et al., 2024)

### sfRNA structure prediction and covariation analysis

Secondary structures 3’ UTR of MVEV were predicted using ClustalW to find xrRNA structure homology within MVEV, followed by folding using LocARNA (http://rna.informatik.uni-freiburg.de/LocARNA/Input.jsp). Pseudoknots were located manually. Secondary structures of 5’ UTR of MVEV were predicted based on previous analysis of the SLA structure (Samanta, 2021), followed by homology analysis using ClustalW. All structures were visualized using VARNA v3-93.

Stockholm formatted alignments of sequences from different genotypes, generated using Clustal Omega (https://www.ebi.ac.uk/jdispatcher/msa/clustalo?stype=rna), and consensus secondary structures were used to build covariance models using R-scape software (http://eddylab.org/R-scape/), using an E-value of 0.05.

### Statistics

Statistical analyses of experimental data were performed using GraphPad Prism 8. The *t* test was used when the difference in variances was <4-fold, skewness was >-2, and kurtosis was <2. Otherwise, the non-parametric Kolmogorov-Smirnov exact test or Mann-Whitney test was used. Paired *t* test was used for comparing neutralisation of different viruses. Kaplan-Meier statistics were determined by log-rank (Mantel-Cox) test.

## Results

### Phylogenetic analysis of MVEV isolates

48 isolates from the Doherty Collection, collected between 1950 and 1980, were found to be labelled as MVEV. After testing for cytopathic effect on BHK cells, complete genomes encoded by 43 MVEV isolates from the Doherty Collection were sequenced. Bioinformatic analyses identified 34 of the sequenced isolates as MVEV. Pairwise identities between isolates starting with “212” calculated using PrME, full genome, and NS1-NS5 sequences shows extremely high levels of genetic identity to each other (> 99.5% nt). Additionally, 1/56_M_1964 was found to be identical to NGPool and to have 99% identity with NG156 (the former being used for ongoing analyses moving forward). 18441C was also found to be identical to 18441C7/8 (the former being used for ongoing analyses moving forward). Therefore, a total number of 15 unique new isolates were identified from the Doherty Collection. Phylogenetic analyses were performed on these isolates, alongside 9 publicly available full length sequences, previously allocated into four genotypes. The phylogenetic tree estimated using the Maximum Likelihood (ML) method on the full length sequences (Figure 1), shows that most of the Doherty Collection MVEV isolates cluster within G1, with a substantial cohort belonging to the G1B sub-lineage. One isolate, 1_56M_1964, belongs to the G3/4 clade. ML analysis on NS1-NS5 nucleotide and amino acid sequences (Supplementary Figure 1-2) shows similar grouping of isolates.

**Figure 1.**
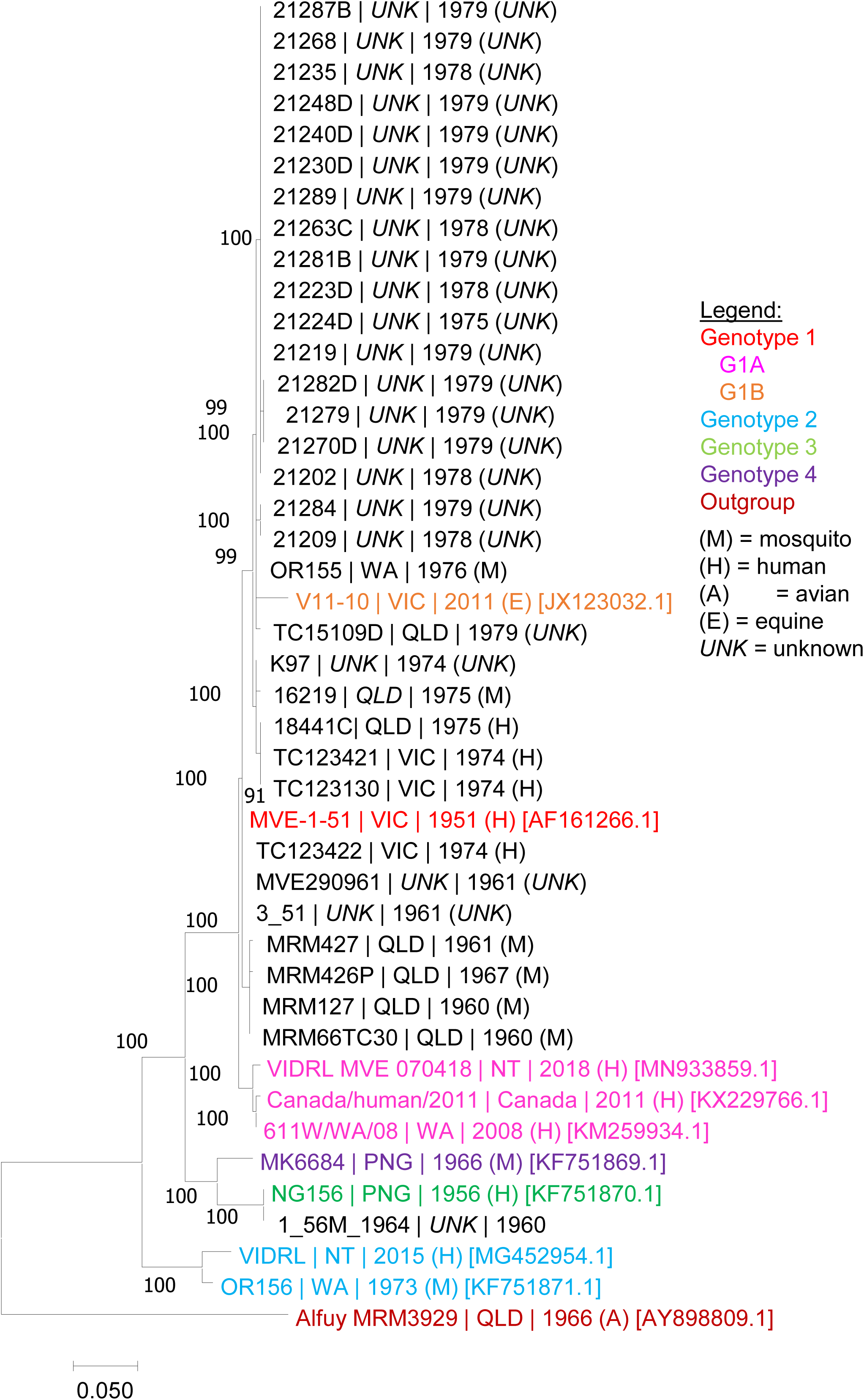
Maximum Likelihood phylogenetic tree estimated using MVEV full genome gene sequences. The phylogenetic tree was constructed using the Maximum Likelihood method and General Time Reversible Model and 1000 bootstrap replicates. The numbers at the nodes represent bootstrap support in percentage of 1000 replicates. The scale bar indicates 0.050 nucleotide substitutions per site. Genotypes and sub-lineages are indicated by colour: Genotype 1, red; Genotype 1A, pink; Genotype 1B, orange; Genotype 2, blue; Genotype 3, green; Genotype 4, purple. Origin of isolation is indicated: H, human; A, Avian; M, mosquito; E, equine. Accession numbers are included in square brackets. The tree was rooted to the phylogenetically related flavivirus, Alfuy MRM3929.

ML analysis on the PrME nucleotide and amino acid sequence was also performed on the same isolates with 53 publicly available PrME sequences of MVEV (Figure 2). Grouping of isolates from the Doherty Collection in this tree is different compared to the full length sequence analysis: apart from 156_M_1964 which still belongs to the G3 clade, all isolates group within G1 and none group in G1B.

**Figure 2.**
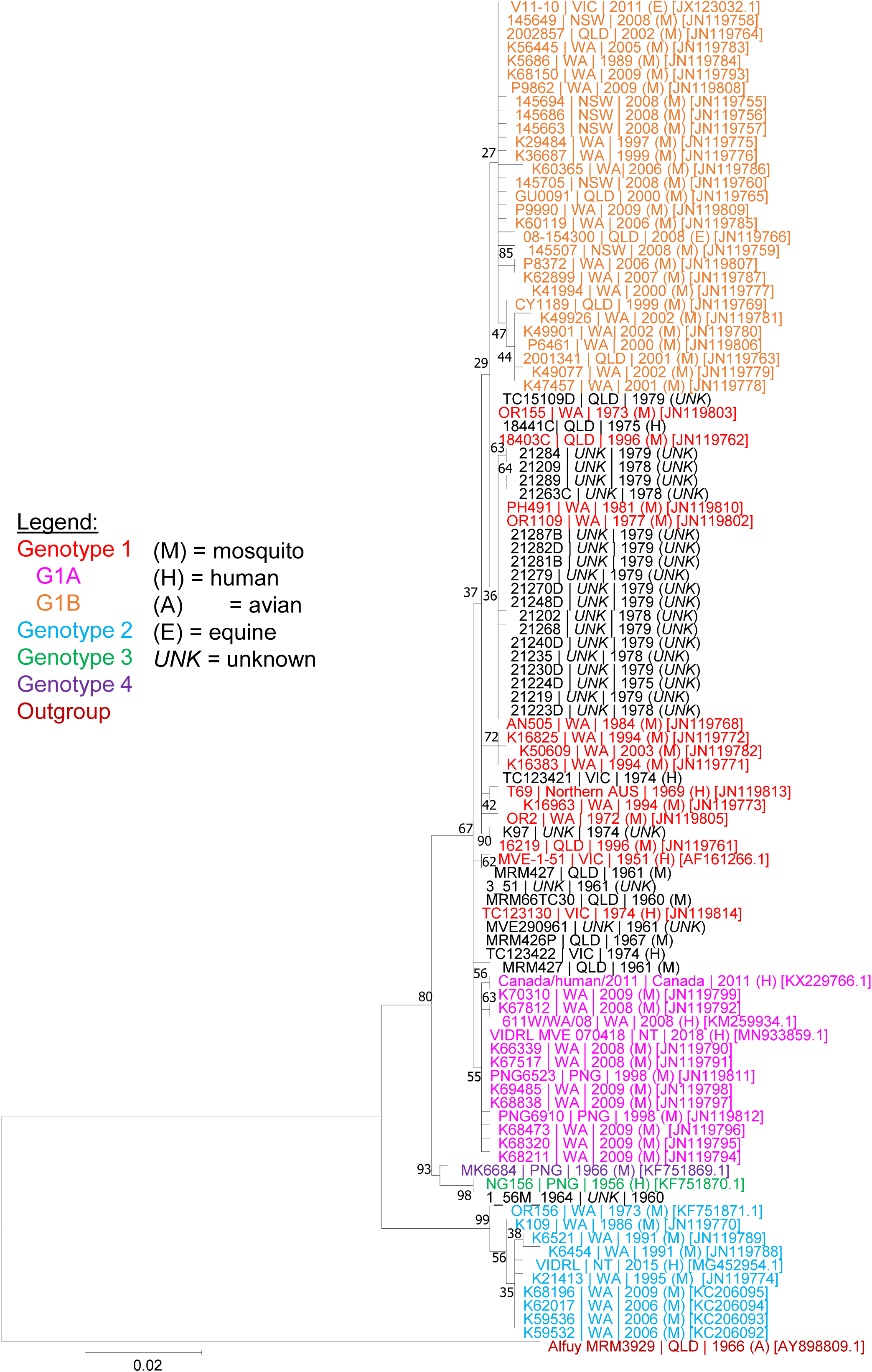
Maximum Likelihood phylogenetic tree estimated using MVEV PrME amino acid sequences. The phylogenetic tree was constructed using the Maximum Likelihood method and General Time Reversible Model and 1000 bootstrap replicates. The numbers at the nodes represent bootstrap support in percentage of 1000 replicates. The scale bar indicates 0.020 amino acid substitutions per site. Genotypes and sub-lineages are indicated by colour: Genotype 1, red; Genotype 1A, pink; Genotype 1B, orange; Genotype 2, blue; Genotype 3, green; Genotype 4, purple. Origin of isolation is indicated: H, human; A, Avian; M, mosquito; E, equine. Accession numbers are included in square brackets. The tree was rooted to the phylogenetically related flavivirus, Alfuy MRM3929.

Overall, the MVEV isolates from the Doherty Collection cluster within G1, with one isolate clustering within G3.

### MVEV Genotype and sub-lineage defining amino acid differences

As previously determined by Williams et al. (2015), each MVEV genotype or sub-lineage has one or more amino acid residues in the PrME region found to be unique to or defining of the isolates within that genotype. Using the publicly available full length genome sequences and the additional full length genome sequences from the Doherty Collection, a number of unique amino acids were identified across prME and the NS regions of MVEV (Table 1). G2 had the most number of unique amino acids, mimicking the genetic analyses demonstrated in the phylogenetic tree with the clustering being the most distant from other genotypes (Figure 1 and 2). G2 had a large number of unique acids, particularly located in the non-structural protein region of the MVEV genome. No unique amino acids were specific to the isolates clustering in G1 that were not part of the G1A or G1B sub-lineage. A number of conservative and non-conservative changes were identified (Supplementary Table 4) across the genotypes.

**Table 1.** Number of unique amino acids in different regions of the MVEV genome that define genotypes or sub-lineages. Unique amino acids are defined as those that are specific to a given genotype and differ from those in other genotypes, which share identical amino acid sequences.

| Sites | G1 | G1A | G1B | G2 | G3 | G4 |
| --- | --- | --- | --- | --- | --- | --- |
| prM | 0 | 0 | 1 | 0 | 1 | 0 |
| E | 0 | 1 | 0 | 14 | 3 | 1 |
| NS1 | 0 | 1 | 2 | 6 | 6 | 1 |
| NS2AB | 0 | 0 | 6 | 7 | 0 | 0 |
| NS3 | 0 | 0 | 1 | 11 | 0 | 0 |
| NS4AB | 0 | 0 | 2 | 14 | 1 | 2 |
| NS5 | 0 | 0 | 3 | 15 | 1 | 3 |

### MVEV infects and replicates in human neuronal REN cells

Human neuronal REN cells were used to determine MVEV neuronal replication capacity in and the subsequent cytopathogenicity. After maturation, the cells were infected with 5 × 10^3^ CCID_50_ of the indicated MVEV isolates, chosen for their respective genotypes. At 7 dpi, microscopy images were taken. All isolates showed cytopathic effect, with OR156 (G2) showing the lowest effect (Figure 3A). Isolates K97 (G1), AN505 (G1), and CY1189 (G1B) showed the highest cytopathic effect (Figure 3A). Viral titres from samples taken from the infected cells every 24 hours for 3 dpi (Figure 3B) confirm these results. All isolates showed viral titres, with OR156 (G2) again showing lower levels than the other isolates.

**Figure 3.**
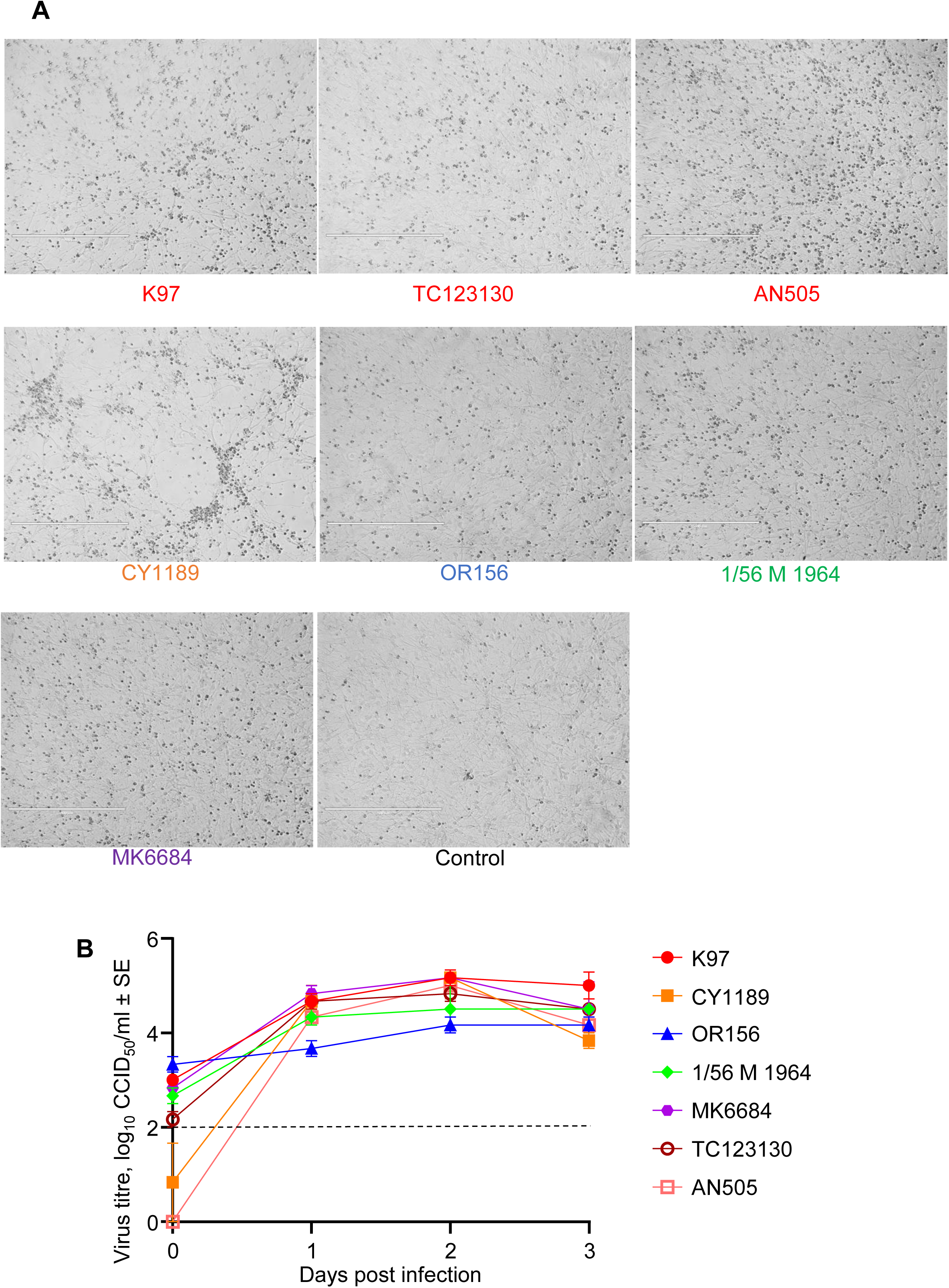
Phenotype of MVEV in mature RENcells. Cells were infected with 5× 10^3^ CCID_50_ of indicated MVEV isolates. **A** Cytopathic effects of tested MVEV isolates in mature RENcells 7 days post infection. **B** Virus titers of the tested MVEV isolates over 3 days. Data shows the mean of n = 3 per group and error bars represent standard error, with limit of detection at 2log_10_CCID_50_.

### C57BL/6J mice infected with MVEV G1, G1B and G4 genotypes led to viral neuroinvasion

Six-week-old adult female C57BL/6J mice were infected s.c. with 5 × 10^5^ CCID_50_ of the aforementioned viruses. Viraemia was similar for K97, 1/56M1964, and MK6684, peaking 2 dpi at 4-5 log_10_CCID_50_/ml of serum. For OR156, viraemia was below the limit of detection of 2 log_10_CCID_50_/ml of serum, and CY1189 did not show any detectable viraemia at all (Figure 4A-E). A qPCR was performed on serum from mice infected with CY1189 to confirm the presence of viral RNA, which came out positive (data not shown). The infection appeared to stall weight gain for most mice until day ∼15 (Figure 4F; orange lines). Between 9 and 14 dpi, four mice (2 x K97, 1 x CY1189, 1 x MK6684) out of a total of 30 mice showed weight loss Η 20% and were euthanized (Figure 4F, †). Kaplan-Meier survival curves provided no significant differences for the viral isolates (Fig. 4G). Euthanized mice also showed varying levels of abnormal posture (hunching), reduced activity, and fur ruffling on the day of euthanasia (data not shown). The four euthanized mice had high levels of brain infection (∼ 8-10 log_10_CCID_50_/mg) and detectable levels of viral titers in the spleen (Fig. 4H). Viral titers in the brain and spleen were only assessed in euthanized mice. Euthanizing healthy mice at corresponding time points was avoided to ensure the generation of a Kaplan-Meier survival curve.

**Figure 4.**
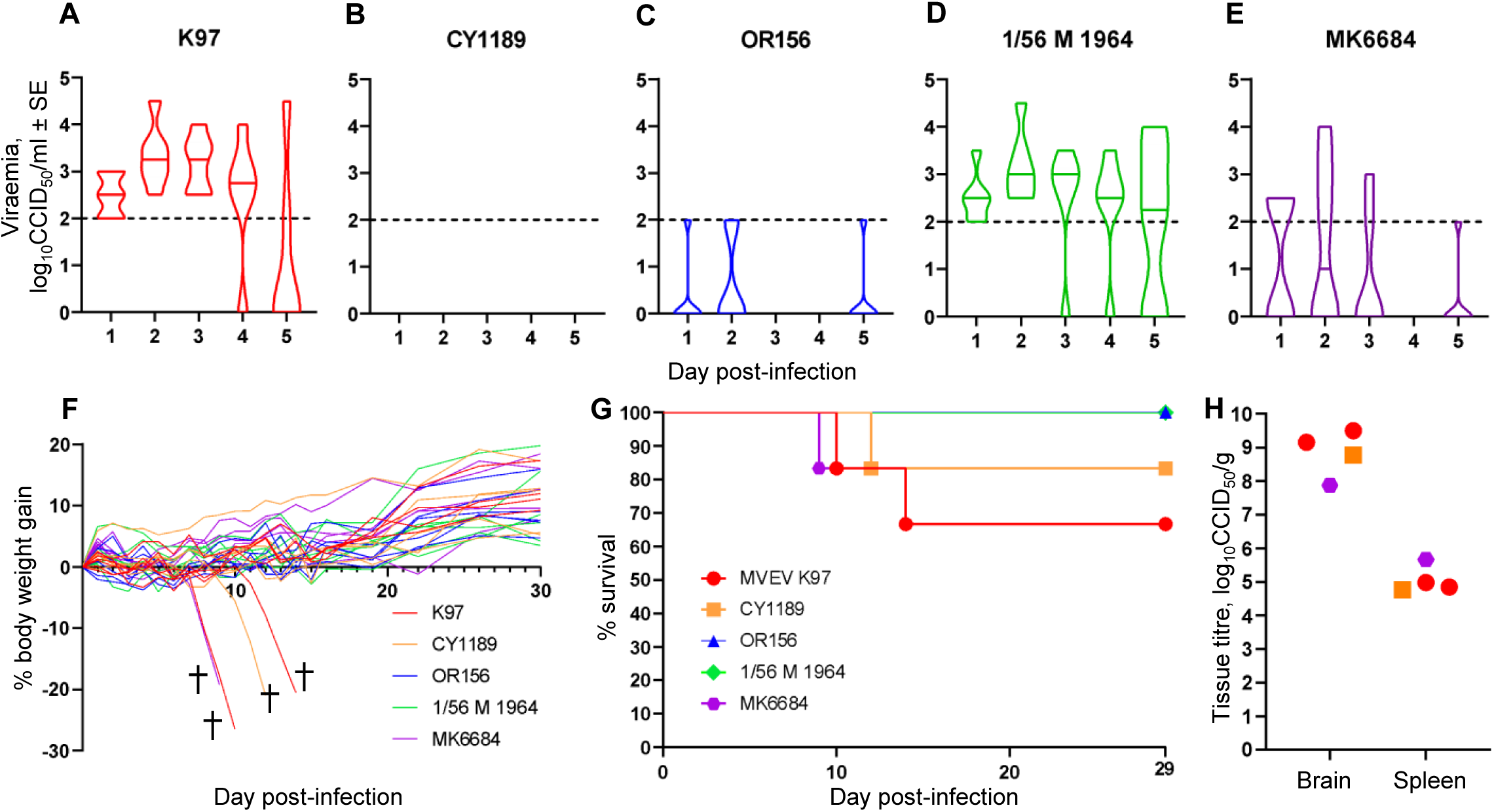
Phenotype of MVEV in C57BL/6J wild-type mice. Female 6 week old C57BL/6J mice were s.c. infected with 5 × 10^3^ CCID_50_ of indicated MVEV isolates. **A-E** Violin plots for n = 6 per group for 5 days are shown for each of the indicated MVEV isolates. Limit of detection is 2 log_10_CCID_50_/ml. **F** Percent body weight change of individual mice after infection with the indicated MVEV isolate at 5 × 10^3^ CCID_50_ compared to individual baseline weight on day zero. Four mice reached ≥20% weight loss and were euthanized (✝). **G** Kaplan-Meier plot showing percent survival (n = 6 for each virus isolate inoculated at 5 × 10^3^ CCID_50_). **H** Viral tissue titers in brain and spleen of four euthanized mice. Tissue titers determined by CCID_50_ assay, with limit of detection at 2 log_10_CCID_50_/g

### MVEV causes severe histological lesions in C57BL/6J mouse brains

H&E staining of MVEV-infected brains revealed a series of histopathological lesions in the mice that succumbed to the virus (Figure 5). Neuronal degeneration, including presence of hyper-eosinophilic and pyknotic nuclei neurons, and neuronal vacuolation (Figure 5A) were identified. Perivascular cuffing (Figure 5B), leukocyte infiltrates (Figure 5C), microgliosis (Figure 5D), and haemorrhagic lesions (Figure 5E) were also identified. These lesions were consistent across all mice that succumbed to MVEV infection. Overall, the observed lesions in the mouse brains are consistent with H&E detectable lesions in post-mortem human MVEV-infected brains.

**Figure 5.**
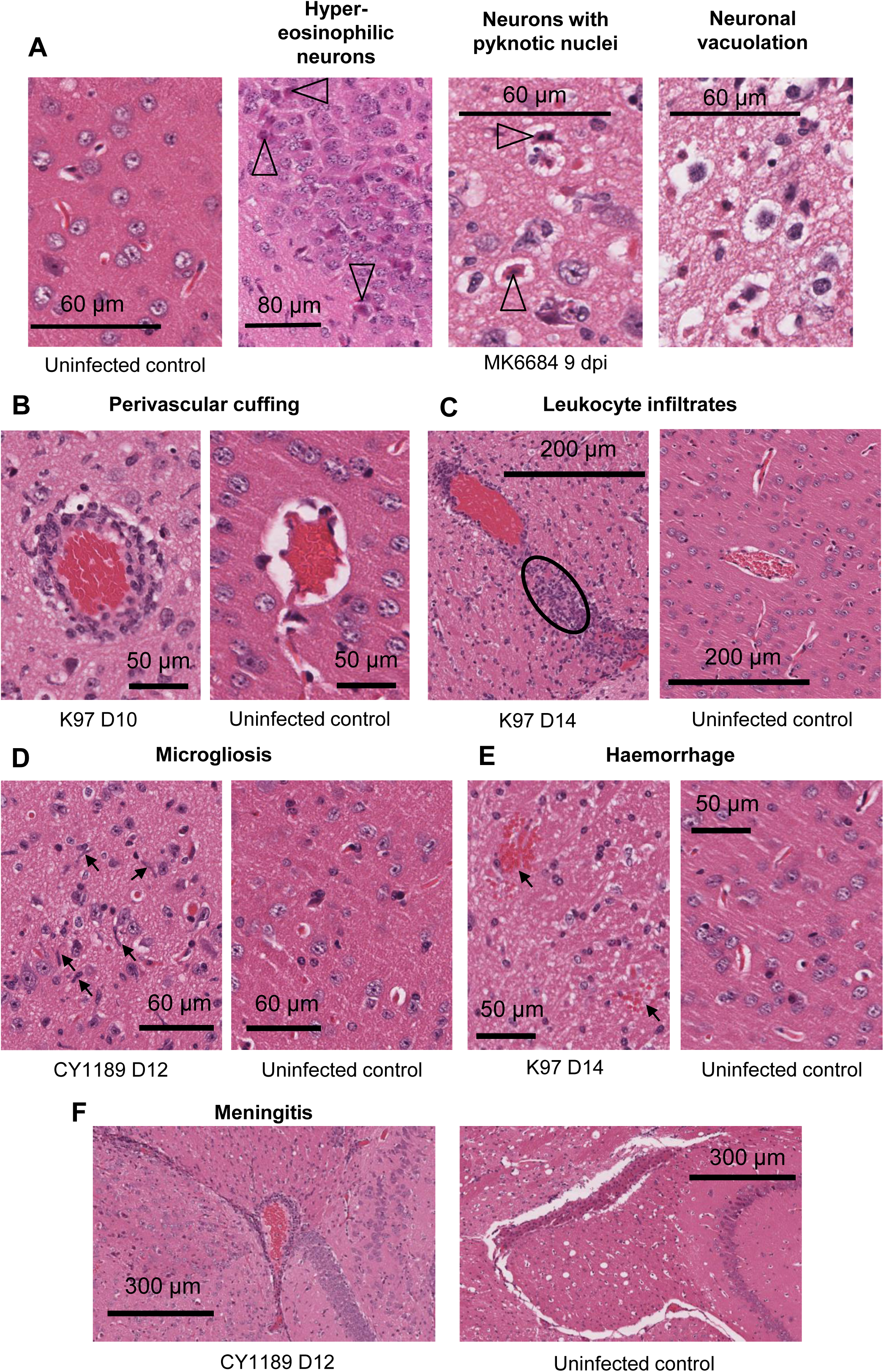
Histological staining and lesions in brains of MVEV-infected mice. MVEV infected brains stained with H&E compared to uninfected control for histological lesions including **A** Neuronal degeneration (indicated by the hyper-eosinophilic neurons, pyknotic nuclei neurons and neuronal vacuolation). Features highlighted by open head triangle **B** Perivascular cuffing indicated by the infiltrates surrounding extravascular vessels **C** Leukocyte infiltrates as highlighted by the circle. **D** Microgliosis is indicated by accumulation of microglia with elongated rod-shaped nuclei (arrows). **E** Haemorrhagic lesions by the arrows. **F** Meningitis as indicated by the infiltrates surrounding the blood vessels in the meninges.

The brains of infected mice were analysed by immunohistochemistry (IHC) using the pan-flavivirus anti-NS1 antibody 4G4 (Figure 6), which detects the viral non-structural protein 1, a highly conserved protein important for flavivirus replication (Rastogi et al., 2016). Staining was consistently seen in the cerebral cortex, hypothalamus, and hippocampus regions of mice that succumbed to the virus, as shown by the representative brain of a K97 infected mouse (Figure 6A). Infection of the cortex and thalamus parallels IHC data from post-mortem human brains infected with the closely related JEV. Staining was not identified in uninfected control mouse brains (Figure 6B).

**Figure 6.**
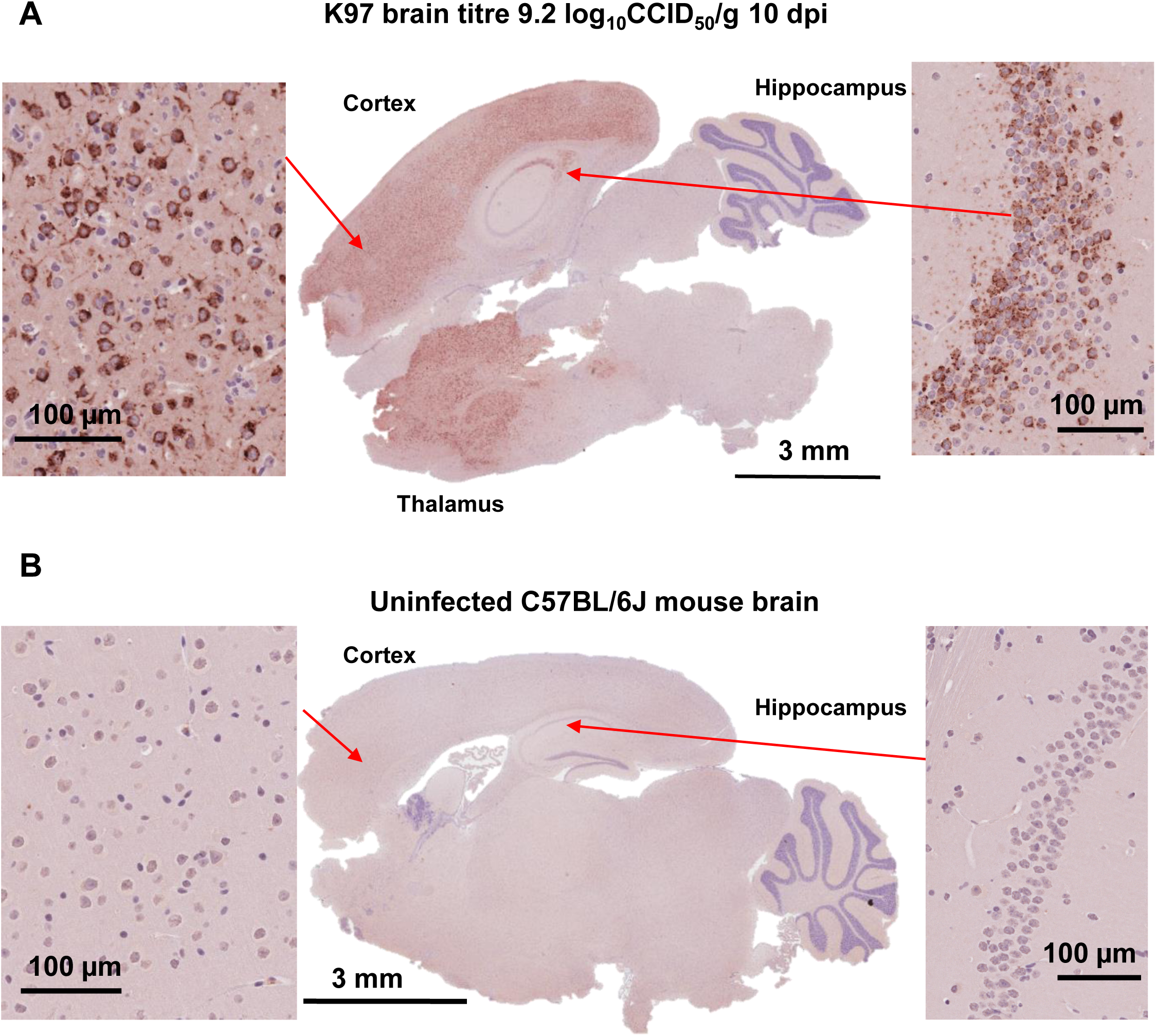
Immunohistochemistry staining in brains of MVEV-infected mice. **A** Immunohistochemistry (IHC) using the pan-flavivirus anti-NS1 antibody 4G4 of a MVEV infected mouse brain (representative shown is a G1 K97 infected brain, brain titre 9.2 log_10_CCID_50_/g. Infected neurons and brain cells indicated in the cortex, hippocampus and thalamus. **B** As for A for an uninfected mouse brain.

### MVEV UTR structure analysis reveals conservation of both 5’ and 3 ’UTR RNA elements

The flaviviral 5’ and 3’ untranslated regions (UTRs) contain RNA elements that are essential for translation, replication, and pathogenesis (Ng et al., 2017). Therefore, 5’ and 3’ UTR consensus structures were generated for MVEV sequences, followed by covariation analysis to determine structural differences between the isolates used in this study.

MVEV 3’ UTR consensus structure was determined using ClustalW and LocARNA, with pseudoknots (PK) located based off phylogenetically related structures from WNV and JEV (Fig. 7A). It contains four stem loops (SL), two dumbbell structures (DB), and a terminal 3’-stem loop (3’SL). SL-II and IV, as well as both DBs, contain PKs. SL-I and its surrounding region has the highest variation in the 3’UTR of MVEV, since OR156 is missing this structure as reported previously (Williams et al., 2014). Additionally, this region shows variation in presence and identity of nucleotides between the other four isolates (Fig. 7B). The other regions are shown to be highly conserved between the different genotypes.

**Figure 7.**
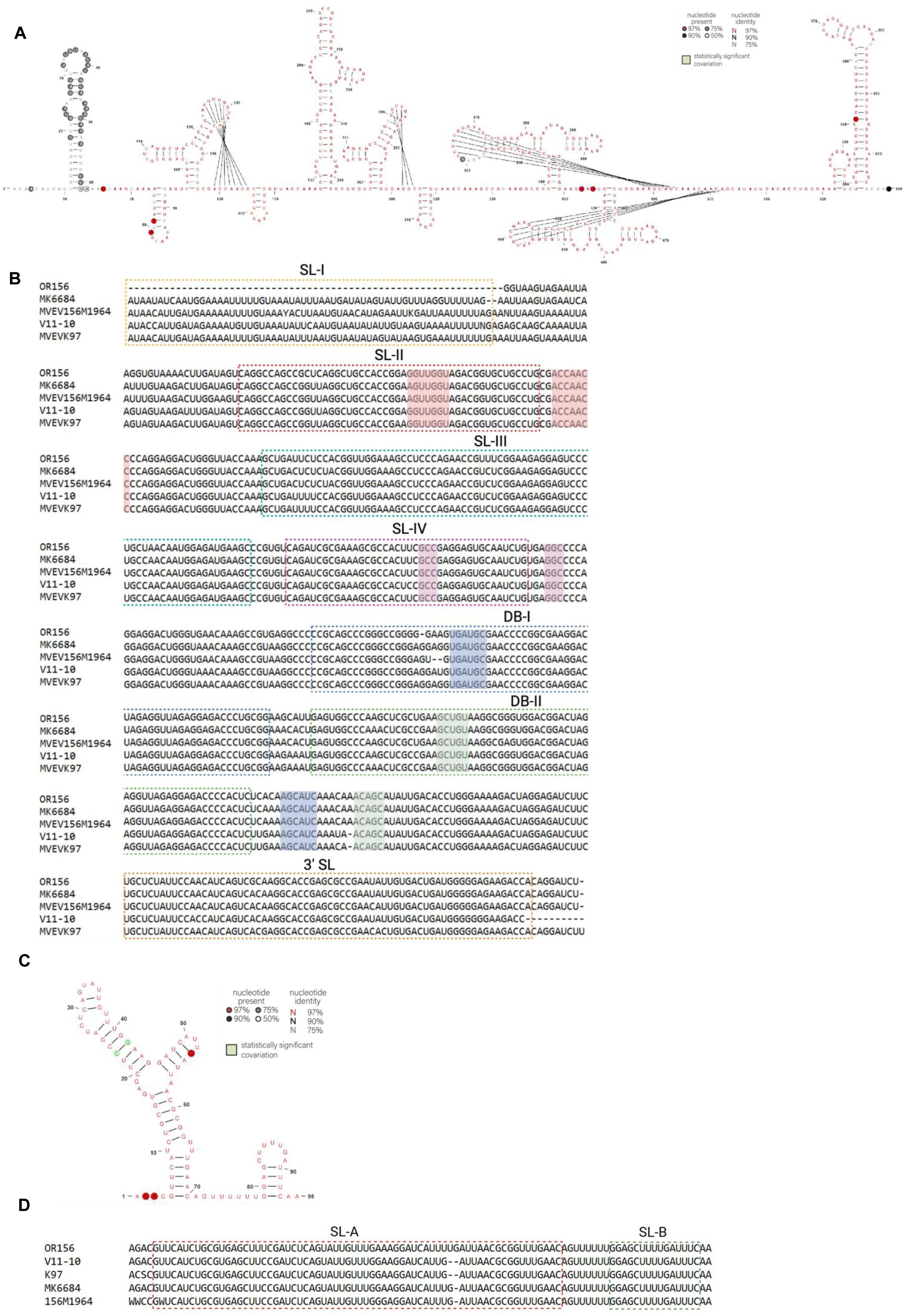
Consensus structures of 5’ and 3’ UTR of MVEV. **A** Consensus structure of the 3’UTR of MVEV based on the covariance model. **B** Sequence alignment of the 3’UTR of MVEV isolates from different genotypes as used in this study. Since no full-length sequence is available for CY1189, it was substituted with V11-10 from G1B. Different structures are indicated by coloured dotted lines, and pseudoknots are indicated by blocks in colours corresponding to its related structures. **C** Consensus structure of the 5’UTR of MVEV based on the covariance model. **D** Sequence alignment of the 5’UTR of MVEV isolates from different genotypes as used in this study. Different structures are indicated by coloured dotted lines.

MVEV 5’ UTR consensus structure was also determined in the same way as the 3’UTR (Fig. 7C). It contains SL-A with a distinct Y-shaped structure, followed by SL-B. The structures are separated by a poly(U) sequence to allow for proper functioning. The whole region is highly conserved in presence of nucleotides (Fig. 7D), and shows statistically significant covariation at locus 23 and 42 in SL-A (Fig. 7C). Overall, both 5’ and 3’ UTR show high conservation between different genotypes of MVEV.

## Discussion

Herein, we provide a comprehensive phylogenetic analysis of full-length sequences of MVEV from the Ralph Doherty collection. Out of the 43 sequenced isolates from the Doherty collection, 15 new unique sequences of MVEV were found after bioinformatic analyses. A majority of these isolates belong to G1, reconfirming previous findings that this is the dominant genotype of the species. No new members of G2 could be added; this lineage is thought to belong to a specific ecological niche, due to the early divergence from a common ancestor. One isolate in G3 was identified, identical to the existing G3 isolate, NG156.

Previous research by Williams et al. (2015) identified two sub-lineages within G1, namely G1A and G1B, based on phylogenetic analyses performed solely on the PrME sequences of MVEV. This study adds an additional 34 full length MVEV sequences (15 of which are unique) for further phylogenetic analyses, also examining differences in the non-structural polyprotein region which has not previously been assessed to the best of our knowledge. The new sequences from the Doherty Collection do not converge the same between full genome nucleotide and PrME amino acid analyses: while new isolates all belong to G1 in the PrME tree, they are divided over G1 and G1B in the full-length tree. These differences are due to the additional genomic data provided by the new full length sequences, and it could mean that the relationship between G1 and G1B is different than previously assumed. However, only one full length sequence of G1B is publicly available. Sequencing of the full length genome of other strains belonging to this genotype in the PrME analysis could further determine this relationship.

In this study, we also use six-week-old adult female C57BL/6J mice to establish models of MVEV infection and disease that recapitulate aspects of human MVEV disease. The capacity of MVEV to cause lethal neuroinvasive infection in mice was four out of 30 infected C57BL/6J mice across 3 different genotypes (G1, G1B and G4). This is slightly lower than what is seen in humans, where a 15-30% case fatality is seen. Mice that succumbed to the virus were infected with isolates K97 (2 mice; G1), CY1189 (1 mouse; G1B), and MK8864 (1 mouse; G4). All of these mice shared similar signs of neuroinvasion. IHC staining using 4G4 antibody and H&E stained histopathology did not show significant differences between brains. Since no IHC or H&E data exists from human MVEV-infected brains, comparisons were made with the closely related JEV. Localisation of IHC stained cells and neuropathological lesions occurred in the cerebral cortex and thalamus, the same regions affected in JEV infection in humans. Additionally, lesions like perivascular cuffing, hemorrhage, and vacuolation are recapitulated in human infection of JEV (German et al., 2006; Iwasaki et al., 1986; Johnson et al., 1985; Maamary et al., 2023; Waller et al., 2022). MRI scans on MVEV patients show bilateral thalamic lesions, thus confirming the thalami are affected in MVEV infection (Derrick et al., 2024; Kienzle and Boyes, 2003; Knox et al., 2012).

Interestingly, three of the mice with fatal neuroinvasion belonged to G1 including subtype G1B, which is also seen as the predominant genotype of MVEV. Previous research has identified Env residue 229, located on the “h”-loop of the DII domain, as a potential driver for higher fitness of G1 (Williams et al., 2015). In other flaviviruses, mutations in this region affected the ability of both group and type-reactive neutralising antibodies to bind (Crill and Chang, 2004; Oliphant et al., 2006), therefore possibly allowing G1 to evade the immune system. Additionally, residue 275, located in the DI-DII hinge region, is variable between genotypes. This region is hypothesised to be involved with viral entry, and mutations to amino acids in this region are shown to affect neuroinvasion and fusion properties of the protein in MVEV and other members of the JEV serocomplex (Hurrelbrink and McMinn, 2001; McMinn et al., 1995; McMinn et al., 1996; Monath et al., 2002). While this research did find several unique amino acids for all genotypes outside of the PrME, additional validation through mutagenesis studies will need to be performed to confirm speculations for contributions to G1 pathogenesis.

The isolate OR156 (G2) showed less virulence compared to the other isolates both *in vitro* and *in vivo*. As part of G2, OR156 belongs to the genotype that diverged early from the other genotypes. Previously, several unique amino acids in the envelope protein for this genotype have been described that could have biological significance (Williams et al., 2015), and it has been previously proposed that these could affect the virulence of members of G2 (Johansen et al., 2007). This study found two unique amino acid changes for G2 in the NS2B and NS3 proteins, at residues Asn84 and Thr132, respectively. These residues belong to the binding pocket of two-component protease NS2B/NS3, responsible for processing the viral polyprotein to the mature viral proteins involved in viral pathogenesis (Yang et al., 2009). Amino acid changes in these same residues between WNV and JEV are shown to cause rearrangement of the possible H-bonds between ligand and enzyme pockets, leading to differences in cleavage efficiency of substrates for the NS2B/NS3 protease (Junaid et al., 2012). Changes in this protease in MVEV could thus also lead to the decreased pathogenicity of OR156 and other members of G2.

This study generated complete consensus structures for both the 5’ and 3’ UTR of MVEV, followed by covariance analyses to determine structural conservation or variation between different genotypes. Overall, nucleotide presence and identity is above 97% for both consensus structures, with the exception of SL-I in the 3’UTR due to the loss of this structure in OR156. Only one statistically significant covariant pair was found in SL-A of the 5’UTR, suggesting this structure is conserved. However, the analyses done in this study can be considered low-power due to the low number of 5’ and 3’ UTR sequences available for all genotypes, which is likely the reason no more covariance was found. While covariance is a stronger indicator of structure conservation of RNA structures, a high conservation of nucleotide identity and presence can also indicate structure conservation (Rivas et al., 2020), as is the case in this study. It is unclear what the effect of the loss of SL-I is on OR156, as the functionality of this structure is currently unknown. While the 3’UTR of related WNV and JEV also contain a similar SL-I (Chapman et al., 2014; Xing et al., 2021; Zhang et al., 2022), some other flaviviruses like Dengue virus do not possess this structure (Villordo et al., 2015). The sequence in which SL-I lays is also degraded by XRN1 before it stalls at SL-II, which has a highly conserved structure amongst flaviviruses (Clarke et al., 2015). It can therefore be hypothesized that SL-I in flaviviruses belonging to the JEV serotype does not have a critical function during infection, if it has a function at all. An RNA mutagenesis study comparing the effect of deletion in a more virulent strain than OR156 could provide more insights.

Due to the current rise in cases of MVEV and the lack of therapeutics, it is vital to monitor the disease and prevent further spread of the disease in alternative ways. Surveillance is an important measure to predict new outbreaks and warn the public as early as possible. Measures like mosquito surveillance and sentinel chickens have already been implemented to monitor MVEV in regions where the virus is prevalent (Braddick et al., 2023; Broom et al., 2001; Campbell and Hore, 1975; Doherty et al., 1976; Victoria Departmet of Health, 2024). To further predict spread, some models have been created (Ho et al., 2016; Selvey et al., 2014b). However, the exact transmission cycle of MVEV is still somewhat unclear. While waterfowl like herons are thought to be an important host (Anderson, 1953), MVEV is also found in other vertebrates like horses, crocodiles, and marsupials (Carver et al., 2009; Gordon et al., 2012; Habarugira et al., 2022; Kay et al., 1985), and their role in the transmission cycle of MVEV remains unclear. More knowledge on the cycle of MVEV and its ecological niche would help predict transmission patterns more accurately. While this study provides new insight in the genetic relationship between genotypes, all added sequences were isolated in the previous century. A significant challenge was presented with the phylogenetic analysis is the lack of records detailing the origins of these viruses. With the exception of a collection date, the passage history, species the viruses were isolated from, and location, applying spatial temporal prediction models to identify regions of risk, potential emerging genotypes and favourable mutations remains a limitation. Addition of contemporary full length sequences can provide insights on the MVEV strains currently circulating, allowing for a more directed approach to producing therapeutics against MVEV.

## Conclusion

In conclusion, 15 new full-length sequences of MVEV were found, providing new insight on the genetic relationship between its genotypes. Additionally, we show that characterisation of MVEV isolates in C57BL/6J mice recapitulates the disease patterns seen in humans, thus establishing a mouse model of disease for MVEV. In the future, this model can be used to study mechanisms of disease and to test novel interventions against the virus.

## Supplementary information

Supplementary Table 1. Established mouse models of MVEV.

Supplementary Table 2. MVEV isolates used in this study.

Supplementary Table 3. Unique amino acids in different regions of the MVEV genome that define genotypes or sub-lineages.

Supplementary Figure 1. Maximum likelihood phylogenetic tree of NS1-NS5 full length nucleotide sequences.

Supplementary Figure 2. Maximum likelihood phylogenetic tree of NS1-NS5 full length amino acid sequences.

