## Supplementary Tables and Figures for "Characterisation and phylogenetic analysis of Murray Valley encephalitis virus (MVEV) from the Ralph Doherty collection"

**Supplementary Table 1. Established mouse models of Murray Valley encephalitis virus (MVEV).**

**Established young MVEV mouse models.**

| Virus isolate/s | Origin | Passage history | Dose/s, route/s, and location/s of inoculation | Mouse strain and age | Disease manifestation | Reference |
|---|---|---|---|---|---|---|
| MVE-F3/51 | Human (?) isolate, Mooroopna 1951 | 3 Tissue culture virus passage | i.p.<br>$10^3$ pfu | BALB/c<br>18-20 days | *Vaccine study*<br>Mortality | Broom, A.K., Wallace, M.J., Mackenzie, J.S., Smith, D.W. and Hall, R.A. (2000), Immunisation with gamma globulin to Murray Valley encephalitis virus and with an inactivated Japanese encephalitis virus vaccine as prophylaxis against Australian encephalitis: Evaluation in a mouse model. J. Med. Virol., 61: 259-265. https://doi.org/10.1002/(SICI)1096-9071(200006)61:2<259::AID-JMV13>3.0.CO;2-M |
| K 12811 | UNK, Kimberley 1993 | 4 Tissue culture virus passage | | | | |
| MVE-1-51 | Human isolate, Mooroopna 1951 | SMB | i.c. or i.p.<br>30 ul<br>0-4 log PFU | Swiss outbred<br>3 weeks | <u>0 log:</u><br>- i.p. 0% mortality<br>- i.c. 0% mortality<br><u>1 log:</u><br>- i.p. 0% mortality<br>- i.c. 50% mortality<br><u>2 log:</u> | Lee E, Lobigs M.2000.Substitutions at the Putative Receptor-Binding Site of an Encephalitic Flavivirus Alter Virulence and Host Cell Tropism and Reveal a Role for Glycosaminoglycans in Entry. J Virol74:.https://doi.org/10.1128/jvi.74.19.8867-8875.2000 |

| | | | | | | |
|---|---|---|---|---|---|---|
| | | | | | - i.p. 55% mortality<br>- i.c. 75% mortality<br><u>3 log:</u><br>- i.p. 100% mortality<br>- i.c. 100% mortality<br><u>4 log:</u><br>- i.p. 100%<br>- i.c. 75%<br><u>5 log</u><br>- i.p. 100%<br>- i.c. 100% | |
| MVE-1-51 | Human isolate, Mooroopna 1951 | SMB | i.c. and i.p.<br>30 ul serial 10-fold dilutions | Outbred Swiss<br>3 weeks | Mortality expressed in LD50, observed over 14 days | M. Lobigs, I.D. Marshall, R.C. Weir, L. Dalgarno. 1988. Murray Valley encephalitis virus field strains from Australia and Papua New Guinea: Studies on the sequence of the major envelope protein gene and virulence for mice. Virology, Volume 165, Issue 1, Pages 245-255. https://doi.org/10.1016/0042-6822(88)90678-2. |
| BH3479 | *Cx. Annulirostris*, Bermah forest 1974 | | | | | |
| TC123130 | Human isolate, Culgoa 1974 | | | | | |
| OR2 | *Cx. Annulirostris*, | | | | | |

| | | | | | | |
|---|---|---|---|---|---|---|
| | Kunnanurra 1972 | | | | | |
| MRM15019 | *Cx. Annulirostris*, Kowanyama 1972 | | | | | |
| MVE-1-56 | Human isolate, Brown river 1956 | | | | | |
| MK6684 | Mixed mosquitoes, Sepik river 1966 | | | | | |
| MVEV-1-51 | Human isolate, Mooroopna 1951 | UNK | i.c.<br>$10^4$ pfu | BALB.A2G-*Mx1* ( B6.A2G-*Mx1* with Mx1-negative BALB/c)<br>3 weeks | Mortality (mice died at 5 days)<br>NS1 staining on three brain sections on different days p.i., number of infected cells counted | Frese M, Lee E, Larena M, Lim PS, Rao S, Matthaei KI, Khromykh A, Ramshaw I, Lobigs M. Internal ribosome entry site-based attenuation of a flavivirus candidate vaccine and evaluation of the effect of beta interferon coexpression on vaccine properties. J Virol. 2014 Feb;88(4):2056-70. doi: 10.1128/JVI.03051-13. Epub 2013 Dec 4. PMID: 24307589; PMCID: PMC3911551. |
| | | | i.p. 1000 pfu | BALB/c<br>23-27 days | Mortality (0% survival at 8 days) | |

| | | | | | | | |
|---|---|---|---|---|---|---|---|
| | | | | 100 pfu | | Mortality (20% survival at 14 days) | |
| | | | | 10 pfu | | Mortality (60% survival at 14 days) | |
| | | | i.c.<br>$10^4$ pfu | | BALB.A2G-*Mx1* ( B6.A2G-*Mx1* with Mx1-negative BALB/c)<br>3 week | Mortality (mice died at 5 days)<br><br>NS1 staining on three brain sections on different days p.i., number of infected cells counted | |
| MVE 1-51 | Human isolate, Mooroopna 1951 | UNK | i.p. or i.c.<br>Tenfold serial dilutions (0.6-60000 pfu) | | Swiss outbred<br>3 weeks | Mortality, not specified | Clark, D. C., Lobigs, M., Lee, E., Howard, M. J., Clark, K., Blitvich, B. J., & Hall, R. A. (2007). In situ reactions of monoclonal antibodies with a viable mutant of Murray Valley encephalitis virus reveal an absence of dimeric NS1 protein. Journal of General Virology, 88(4), 1175–1183. https://doi.org/10.1099/vir.0.82609-0 |

| | | | | | | |
|---|---|---|---|---|---|---|
| MVE-F3/51 | Human (?) isolate, Mooroopna 1951 | UNK | i.p. $10^3$ p.f.u. | BALB/c 18-20 days | *Vaccine study* Mortality (29% ‘background’ mortality) | Wallace, M. J., Smith, D. W., Broom, A. K., Mackenzie, J. S., Hall, R. A., Shellam, G. R., & McMinn, P. C. (2003). Antibody-dependent enhancement of Murray Valley encephalitis virus virulence in mice. Journal of General Virology, 84(7), 1723–1728. https://doi.org/10.1099/vir.0.18980-0 |
| MVE 1-51 | Human isolate, Mooroopna 1951 | PSEK | i.p., range of doses (UI) determined by TCID50 | Swiss outbred 3 weeks | Mortality, culled at appearance of first signs of encephalitis Seroconversion | Prow, N. A., May, F. J., Westlake, D. J., Hurrelbrink, R. J., Biron, R. M., Leung, J. Y., McMinn, P. C., Clark, D. C., Mackenzie, J. S., Lobigs, M., Khromykh, A. A., & Hall, R. A. (2011). Determinants of attenuation in the envelope protein of the flavivirus Alfuy. Journal of General Virology, 92(10), 2286–2296. https://doi.org/10.1099/vir.0.034793-0 |
| MVE K68838 | Mosquito isolate, Western Australia 2012 | C6/36 | i.p. 10 $TCID_{50}$ | Swiss CD-1 (mixed gender) 18 days | Mortality (10% survival by day 20)<br><br>Neuropathology score (encephalitis, meningitis, myelitis)<br><br>Cardiac inflammation | Suen, W. W., Prow, N. A., Setoh, Y. X., Hall, R. A., & Helle Bielefeldt-Ohmann. (2016). End-point disease investigation for virus strains of intermediate virulence as illustrated by flavivirus infections. Journal of General Virology, 97(2), 366–377. https://doi.org/10.1099/jgv.0.000356 |

| | | | | | |
|---|---|---|---|---|---|
| | | | i.p. 1000TCID$_{50}$ | | Mortality (10% survival by day 20)<br><br>Neuropatholog y score (encephalitis, meningitis, myelitis)<br><br>Necrosis and mineralisation in cerebral cortex<br><br>Granulomatous inflammation<br><br>Cardiac inflammation |
| MVE 1-51 | Human isolate, Mooroopna 1951 | As described in Lobigs et al., 1986 | i.p. 10TCID$_{50}$ | | Mortality (0% survival by day 7)<br><br>Neuropatholog y score (encephalitis, meningitis, myelitis)<br><br>Cardiac inflammation |

| | | | | | |
|---|---|---|---|---|---|
| | | | i.p.<br>1000TCID$_{50}$ | | Mortality (0% survival by day 8)<br><br>Neuropathology score (encephalitis, meningitis, myelitis)<br><br>Cardiac inflammation |

**Established adult MVEV mouse models.**

| Virus isolate/s | Origin | Passage history | Dose/s, route/s, and location/s of inoculation | Mouse strain and age | Disease manifestations | Reference/s |
|---|---|---|---|---|---|---|
| MVE 1-51 | Human isolate, Mooroopna 1951 | UNK | i.v.<br><br>$10^2$ pfu | IFN-a/b-R $^{-/-}$<br><br>6 weeks | Mortality (0% survival after 6 days) | Clark, D. C., Lobigs, M., Lee, E., Howard, M. J., Clark, K., Blitvich, B. J., & Hall, R. A. (2007). In situ reactions of monoclonal antibodies with a viable mutant of Murray Valley encephalitis virus reveal an absence of dimeric NS1 protein. *Journal of General Virology*, *88*(4), 1175–1183. https://doi.org/10.1099/vir.0.82609-0 |
| MVE-1-51 | Human isolate, Mooroopna 1951 | SMB | i.v. or i.p. ? (methods and results not the same)<br><br>$10^8$ PFU | C57BL/6<br><br>14 weeks | *Vaccine study* Mortality (40-70% survival after 22 days, experiment done twice with different results) | Lobigs M, Larena M, Alsharifi M, Lee E, Pavy M. Live chimeric and inactivated Japanese encephalitis virus vaccines differ in their cross-protective values against Murray Valley encephalitis virus. J Virol. 2009 Mar;83(6):2436-45. doi: 10.1128/JVI.02273-08. Epub 2008 Dec 24. PMID: 19109382; PMCID: PMC2648276. |
| | | | i.v. or i.p.<br><br>$10^2$ to $10^3$ PFU | IFN-α-R$^{-/-}$<br><br>14 weeks | *Vaccine study* Mortality (0% survival at day 9) | |
| | | | i.v. or i.p.<br><br>$10^2$ to $10^3$ PFU | IFN-α/γ-R$^{-/-}$<br><br>14 weeks | *Vaccine study* Mortality (0% survival at day 6) | |
| | | UNK | i.p. | IFNAR$^{-/-}$ | Mortality (0% survival at 6 days) | |

<table>
<tr><td rowspan="4">MVEV-1-51</td><td rowspan="4">Human isolate, Mooroopna 1951</td><td rowspan="4"></td><td colspan="2">$10^{2}$ pfu</td><td>6-8 weeks</td><td></td><td rowspan="4">Frese M, Lee E, Larena M, Lim PS, Rao S, Matthaei KI, Khromykh A, Ramshaw I, Lobigs M. Internal ribosome entry site-based attenuation of a flavivirus candidate vaccine and evaluation of the effect of beta interferon coexpression on vaccine properties. J Virol. 2014 Feb;88(4):2056-70. doi: 10.1128/JVI.03051-13. Epub 2013 Dec 4. PMID: 24307589; PMCID: PMC3911551.</td></tr>
<tr><td rowspan="2">i.v.</td><td>$10^{3}$ pfu</td><td rowspan="2">NOD-scid<br>6-7 weeks</td><td>Mortality (0% survival at 12 days)</td></tr>
<tr><td>$10^{5}$ pfu</td><td>Mortality (0% survival at 9 days)</td></tr>
<tr><td colspan="2">i.c.<br>$10^{4}$ pfu</td><td>BALB.A2G-Mx1 ( B6.A2G-Mx1 with Mx1-negative BALB/c)<br>“adult”</td><td>NS1 or Mx1 staining in Cortex, Olfactory bulb, and ventral striatum, visualized and quantified</td></tr>
<tr><td>MVEV-1-51</td><td>Human isolate, Mooroopna 1951</td><td>SMB</td><td colspan="2">i.v. or i.c.<br>$10^{2}$, $10^{5}$, or $10^{8}$ pfu</td><td>B6<br>6 weeks</td><td><u>$10^{2}$ pfu i.v.</u><br>Mortality (survival 50% at 21 days)<br><u>$10^{5}$ pfu i.v.</u><br>Mortality (survival 60% at 16 days)<br><u>$10^{8}$ pfu i.v.</u><br>- Mortality (survival 0% at 6 days)<br>- Viral pfu in spleen, brain, and muscle (not specified) at 2, 3, or 4 days p.i.<br><u>$10^{2}$ pfu i.c.</u><br>Mortality (survival 0% at 7 days)</td><td>Licon Luna RM, Lee E, Müllbacher A, Blanden RV, Langman R, Lobigs M. Lack of both Fas ligand and perforin protects from flavivirus-mediated encephalitis in mice. J Virol. 2002 Apr;76(7):3202-11. doi: 10.1128/jvi.76.7.3202-3211.2002. PMID: 11884544; PMCID: PMC136025.</td></tr>
</table>

| | | | | | | |
|---|---|---|---|---|---|---|
| | | | | 10 weeks | 10$^8$ pfu i.v.<br>Mortality (survival 20% at 16 days) | |
| | | | | B6 *perf*$^{-/-}$× *gld*<br>6 weeks | 10$^2$ pfu i.v.<br>Mortality (survival 100% at 21 days)<br>10$^5$ pfu i.v.<br>Mortality (survival 100% at 16 days)<br>10$^8$ pfu i.v.<br>- Mortality (survival 0% at 12 days)<br>- Viral pfu in spleen, brain, and muscle (not specified) at 2, 3, or 4 days p.i.<br>10$^2$ pfu i.c.<br>Mortality (survival 0% at 7 days) | |
| | | | | 10 weeks | 10$^8$ pfu i.v.<br>Mortality (survival 25% at 16 days) | |
| | | | | B6 *perf*$^{+/-}$ × *gld*<br>6 weeks | 10$^2$ pfu i.v.<br>Mortality (survival 65% at 21 days) | |
| MVE-1-51 | Human isolate, Mooroopna 1951 | UNK | i.v.<br>10$^2$ pfu | B6<br>6 weeks | - Mortality (70% survival at 21 days)<br>- Brain cell infiltration and necrosis | Lobigs M, Müllbacher A, Wang Y, Pavy M, Lee E. Role of type I and type II interferon responses in recovery from infection with an encephalitic flavivirus. J Gen Virol. |

| | | | | | | |
|---|---|---|---|---|---|---|
| | | | | | - Brain histology:<br>- Perivascular oedema<br>- Vascular congestion<br>- Parenchymal leukocyte infiltration<br>- Meningeal inflammation | 2003 Mar;84(Pt 3):567-572. doi: 10.1099/vir.0.18654-0. PMID: 12604807. |
| | | | | IFN-γ$^{-/-}$<br>6 weeks | Mortality (40% survival at 21 days)<br><br>Brain cell infiltration and necrosis<br><br>Brain histology:<br>- Perivascular oedema<br>- Vascular congestion<br>- Parenchymal leukocyte infiltration<br>- Meningeal inflammation | |
| | | | | IFN-α-R$^{-/-}$<br>6 weeks | Mortality (0% survival at 6 days) | |
| | | | | i-NOS$^{+/+}$<br>6 weeks | Mortality (65% survival at 21 days)<br><br>Brain cell infiltration and necrosis | |

| | | | | | | |
|---|---|---|---|---|---|---|
| | | | | i-NOS$^{-/-}$<br>6 weeks | Mortality (40% survival at 21 days)<br><br>Brain cell infiltration and necrosis<br><br>Brain histology:<br>- Perivascular oedema<br>- Vascular congestion<br>- Parenchymal leukocyte infiltration<br>- Meningeal inflammation | |
| F/3/51 | Human (?) isolate, Mooroopna 1951 | V | i.c.<br><br>(1000-0.1) $TCID_{60}$/5 ul | BALB/c<br><br>10 weeks? | Mortality:<br>- 1000: 0% survival, death at 7-9 days<br>- 100: 10% survival, death at 8-11 days<br>- 10: 50% survival, death at 8-13 days<br>- 1: 100% survival<br>- 0.1: 100% survival | Hall RA, Brand TN, Lobigs M, Sangster MY, Howard MJ, Mackenzie JS. Protective immune responses to the E and NS1 proteins of Murray Valley encephalitis virus in hybrids of flavivirus-resistant mice. J Gen Virol. 1996 Jun;77 ( Pt 6):1287-94. doi: 10.1099/0022-1317-77-6-1287. PMID: 8683218. |
| | | | | C3H/RV<br><br>10 weeks? | Mortality:<br>- 1000: 40% survival, death at 10-14 days<br>- 100: 70% survival, death at 10-12 days<br>- 10: 90% survival, death at 12 days<br>- 1: 100% survival | |

| | | | | | |
|---|---|---|---|---|---|
| | | | | | - 0.1: 100% survival |
| | | | | (BALB/c × C3H/RV) $F_t$<br><br>10 weeks? | <u>Mortality:</u><br>- 1000: 10% survival, death at 7-9 days<br>- 100: 36% survival, death at 11-16 days<br>- 10: 42% survival, death at 9-14 days<br>- 1: 100% survival<br>- 0.1: 100% survival |

**Supplementary Table 2. List of Murray Valley encephalitis virus isolates identified in the Doherty collection.**

| | Isolate | Date | Cytopathic effect on BHK cells | Sequenced |
|---|---|---|---|---|
| 1 | MRM 66 T.C. #30 | 19/09/1900 | ☑ | ☑ |
| 2 | NG Pool | 07/12/1967 | ☑ | ☑ |
| 3 | 1/56 M 1964 | 27/09/1960 | ☑ | ☑ |
| 4 | 3/51 | 09/06/1961 | ☑ | ☑ |
| 5 | MRM 127 | 27/11/1960 | ☑ | ☑ |
| 6 | MRM 74 | 01/07/1960 | ☑ | ☑ |
| 7 | MRM 75 | 08/07/1960 | ☑ | ☑ |
| 8 | MRM 77 | 19/07/1960 | NO | NO |
| 9 | MRM 453 | 07/09/1961 | NO | NO |
| 10 | MRM 427 | 05/08/1961 | ☑ | ☑ |
| 11 | MRM 426P | 05/12/1967 | ☑ | ☑ |
| 12 | MRM 442 | 28/08/1961 | ☑ | ☑ |
| 13 | K 97 | 22/07/1974 | ☑ | ☑ |
| 14 | TC123421 | 26/04/1974 | ☑ | ☑ |
| 15 | 16129 H | 16/09/1974 | ☑ | ☑ |
| 16 | CSIRO 1 (L) | 20/11/1975 | ☑ | ☑ |
| 17 | 16219 | 04/04/1975 | ☑ | ☑ |
| 18 | 21224D | 19/01/1979 | ☑ | ☑ |
| 19 | 18441C | 22/05/1975 | ☑ | ☑ |
| 20 | 21240D | 20/01/1979 | ☑ | ☑ |
| 21 | 21230D | 19/01/1979 | ☑ | ☑ |
| 22 | 21268 | 05/02/1979 | ☑ | ☑ |
| 23 | 21284 | 02/02/1979 | ☑ | ☑ |
| 24 | 21202 | 14/12/1978 | ☑ | ☑ |
| 25 | 21209 | 17/12/1978 | ☑ | ☑ |
| 26 | 21218 | 17/01/1979 | NO | NO |
| 27 | 21223 | 17/12/1978 | ☑ | ☑ |
| 28 | 21235 | 20/12/1978 | ☑ | ☑ |
| 29 | 21263C | 02/02/1979 | ☑ | ☑ |
| 30 | 21267C | 18/02/1979 | NO | NO |
| 31 | 21270D | 05/02/1979 | ☑ | ☑ |
| 32 | 21279 | 02/02/1979 | ☑ | ☑ |
| 33 | 21282D | 05/02/1979 | ☑ | ☑ |
| 34 | 21289 | 02/02/1979 | ☑ | ☑ |
| 35 | MRM472 | 07/09/1961 | ☑ | ☑ |
| 36 | 21219 | 04/01/1979 | ☑ | ☑ |
| 37 | 21248D | 27/01/1979 | ☑ | ☑ |
| 38 | 21281B | 01/02/1979 | ☑ | ☑ |
| 39 | 21287B | 02/02/1979 | ☑ | ☑ |
| 40 | 18403 D1&2 | 18/07/1975 | ☑ | ☑ |
| 41 | 18441 C7&8 | 14/07/1975 | ☑ | ☑ |
| 42 | 18444 C7&8 | 11/07/1975 | ☑ | ☑ |
| 43 | OR155 | 01/01/1976 | ☑ | ☑ |
| 44 | OR156 | 01/01/1976 | NO | NO |
| 45 | MVE | 29/09/1961 | ☑ | ☑ |
| 46 | TC123130 | 07/02/1980 | ☑ | ☑ |
| 47 | TC15109D | 12/04/1979 | ☑ | ☑ |
| 48 | TC123422 | 26/04/1974 | ☑ | ☑ |

**Supplementary Table 3. CCID50 titrations of MVEV stocks.** Log10 CCID50 was calculated at 5 and 7 days post infection, after titration directly onto Vero E6 cells or passaged from C6/36 to Vero E6. New stocks from isolates obtained from UQ were grown onto C6/36 or PK15 cells. as indicated.

| | **D5** | | **D7** | |
|---|---|---|---|---|
| **MVEV strain** | **C6/36 to VERO E6** | **VERO E6** | **C6/36 to VERO E6** | **VERO E6** |
| 18441C | 8.7 | 9 | 8.7 | 9 |
| 18441C78 | 8 | 8.3 | 8 | 8.3 |
| OR156 | 0 | 2.9 | 0 | 2.9 |
| MRM426P | 9 | 8 | 9 | 8 |
| 21219 NEW | 6.8 | 6.7 | 6.8 | 6.7 |
| MRM127 | 8.6 | 7.7 | 8.6 | 7.7 |
| 3/51 | 5.5 | 5.1 | 5.5 | 5.1 |
| 16219 | 9.1 | 8.7 | 9.1 | 8.7 |
| 21202 | 7.8 | 8.4 | 7.8 | 8.4 |
| 21209 | 8.3 | 8.1 | 8.3 | 8.1 |
| 21223 | 9.1 | 8.5 | 9.1 | 8.5 |
| 21235 | 8.8 | 8.7 | 8.8 | 8.7 |
| 21268 | 8.3 | 8.3 | 8.3 | 8.3 |
| 21279 | 7.9 | 7.7 | 7.9 | 7.7 |
| 21284 | 8.1 | 7.1 | 8.1 | 7.1 |
| 21289 | 8.6 | 8.4 | 8.6 | 8.4 |
| 156M1964 | 8.5 | 8.2 | 8.5 | 8.2 |
| 21224D | 5.9 | 4.9 | 5.9 | 4.9 |
| 21230D | 7.6 | 7.3 | 7.6 | 7.3 |
| 21240D | 8.5 | 8.3 | 8.5 | 8.3 |
| 21248D | 8.9 | 8.2 | 8.9 | 8.2 |
| 21263C | 8.4 | 8.1 | 8.4 | 8.1 |
| 21270D | 8 | 8.1 | 8 | 8.1 |
| 21281B | 8.8 | 8.4 | 8.8 | 8.4 |
| 21282D | 8.2 | 7.9 | 8.2 | 7.9 |
| 21287B | 7.3 | 7.7 | 7.3 | 7.7 |

| | | | | |
|---|---|---|---|---|
| K97 | 8.8 | 8.5 | 8.8 | 8.5 |
| MRM 66 TC30 | 7.5 | 7.4 | 7.5 | 7.4 |
| MRM427 | 8.5 | 8.2 | 8.5 | 8.2 |
| MVE 29/09/61 | 7.2 | 7.3 | 7.2 | 7.3 |
| NG Pool | 8.8 | 9.3 | 8.8 | 9.3 |
| OR155 | 6.1 | 6.2 | 6.1 | 6.2 |
| TC123421 | 7.8 | 7.8 | 7.8 | 7.8 |
| NG 156 C6/36 | 7.4 | 6.9 | 7.4 | 6.9 |
| Alfuy C6/36 | 8.3 | 4.9 | 8.3 | 4.9 |
| NG 6684 C6/36 | 6.9 | 7.2 | 6.9 | 7.2 |
| MVE 1-51 C6/36 | 7.5 | 7.4 | 7.5 | 7.4 |
| OR156 C6/36 | 5.9 | 4.6 | 5.9 | 4.6 |
| MVE 1189 C6/36 | 6.5 | 6.4 | 6.5 | 6.4 |
| K30702 C6/36 | 6.2 | 7 | 6.2 | 7 |
| AN 505 C6/36 | 6.1 | 5.9 | 6.1 | 5.9 |
| 18403C C6/36 | 6.9 | 6.9 | 6.9 | 6.9 |
| T69 C6/36 | 6.7 | 7.2 | 6.7 | 7.2 |
| OR2 C6/36 | 8.1 | 7.3 | 8.1 | 7.3 |
| T69 PK15 | 7.9 | 7.1 | 7.9 | 7.1 |
| Alfuy PK15 | 5.2 | 4.6 | 5.2 | 4.6 |
| OR156 PK15 | 7 | 5.8 | 7 | 5.8 |
| NG 6684 PK15 | 8.3 | 8.3 | 8.3 | 8.3 |
| 1189 PK15 | 7.6 | 6.9 | 7.6 | 6.9 |
| OR2 PK15 | 7.8 | 7.4 | 7.8 | 7.4 |
| NG 156 PK15 | 7.8 | 7.5 | 7.8 | 7.5 |
| AN 505 PK15 | 6.5 | 6.9 | 6.5 | 6.9 |

**Supplementary Table 4. Unique amino acids in different regions of the MVEV genome that define genotypes or sub-lineages.** Unique amino acids are defined as those that are specific to a given genotype and differ from those in other genotypes, which share identical amino acid sequences. The location within each site of the genome is given with the amino acid substitution type. Out = outgroup virus (Alfuy).

| Site | Genotype or sub-lineage | | | | | | | Location | Substitution type |
|---|---|---|---|---|---|---|---|---|---|
| | G1 | G1A | G1B | G2 | G3 | G4 | Out | | |
| **Pre-membrane** | | | | | | | | | |
| 24 | A | A | V | A | A | A | T | c strand | Conservative |
| 77 | N | N | N | N | H | N | S | g strand | Non-conservative |
| **Envelope** | | | | | | | | | |
| 15 | A | A | A | V | A | A | V | DI $A_O$-$B_O$ loop | Conservative |
| 21 | V | V | V | I | V | V | V | DI $A_O$-$B_O$ loop | Conservative |
| 55 | L | L | L | V | L | L | V | DII a strand | Conservative |
| 64 | T | T | T | T | T | I | T | DII a-b loop | Non-conservative |
| 72 | S | S | S | A | S | S | S | DII b strand | Non-conservative |
| 126 | A | A | A | T | A | A | S | DII e strand | Non-conservative |
| 157 | T | T | T | S | T | T | T | DI αA helix | Conservative |
| 180 | M | L | M | M | M | M | L | DI $G_O$-$H_O$ loop | Conservative |
| 187 | T | T | T | T | A | T | T | DI $H_O$ strand | Non-conservative |
| 205 | T | T | T | S | T | T | T | DII f strand | Conservative |
| 232 | E | E | E | D | E | E | S | DII i loop | Conservative |
| 270 | I | I | I | I | V | I | I | DII k loop | Conservative |
| 276 | S | S | S | G | S | S | S | | Conservative |
| 330 | T | T | T | A | T | T | L | | Non-conservative |
| 352 | V | V | V | I | V | V | V | | Conservative |
| 369 | A | A | A | S | A | A | S | | Non-conservative |
| 442 | V | V | V | V | I | V | M | | Conservative |
| 461 | S | S | S | T | S | S | T | | Conservative |
| 474 | V | V | V | I | V | V | V | | Conservative |
| **Non-structural protein 1** | | | | | | | | | |
| 21 | I | I | I | V | I | I | V | B roll | Conservative |
| 29 | I | I | I | V | I | I | V | B roll | Conservative |

| **Site** | G1 | G1A | G1B | G2 | G3 | G4 | Out | **Location** | **Substitution type** |
|---|---|---|---|---|---|---|---|---|---|
| **Non-structural protein 1 (continued)** | | | | | | | | | |
| 79 | L | L | L | L | F | L | L | Wing | Conservative |
| 92 | K | K | K | K | R | K | K | Wing | Conservative |
| 112 | D | E | D | D | D | D | E | Wing | Conservative |
| 123 | L | L | F | L | L | L | I | Wing | Conservative |
| 135 | V | V | V | V | I | V | I | Wing | Conservative |
| 176 | T | T | T | S | T | T | T | Wing | Conservative |
| 192 | H | H | H | Y | H | H | L | Central B ladder | Non-conservative |
| 205 | G | G | G | G | E | G | S | Central B ladder | Non-conservative |
| 219 | E | E | E | E | D | E | E | Central B ladder | Conservative |
| 261 | K | K | R | K | K | K | K | Central B ladder | Conservative |
| 271 | E | E | E | D | E | E | E | Central B ladder | Conservative |
| 293 | K | K | K | R | K | K | K | Central B ladder | Conservative |
| 338 | M | M | M | M | V | M | M | Central B ladder | Conservative |
| 353 | F | F | F | F | F | L | L | Central B ladder | Conservative |
| **Non-structural protein 2A** | | | | | | | | | |
| 37 | A | A | A | T | A | A | V | pTMS2 | Non-conservative |
| 98 | R | R | R | K | R | R | H | Loop between pTMS3 and 4 | Conservative |
| 150 | I | I | I | M | I | I | L | pTSM6 | Conservative |
| 177 | I | I | I | V | I | I | I | pTSM7 | Conservative |
| 183 | L | L | M | L | L | L | L | pTSM7 | Conservative |
| 190 | V | V | A | V | V | V | A | pTSM7 | Conservative |
| **Non-structural protein 2B** | | | | | | | | | |
| 31 | V | V | I | V | V | V | I | TMD2 | Conservative |

| | | | | | | | | | |
|---|---|---|---|---|---|---|---|---|---|
| 45 | I | I | V | I | I | I | I | TMD2 | Conservative |
| 65 | G | G | G | E | G | G | D | Loop between TMD2 and 3 | Non-conservative |
| 84 | D | D | D | N | D | D | D | Loop between TMD2 and 3 | Non-conservative |
| 99 | V | V | V | I | V | V | L | TMD3 | Conservative |
| 107 | V | V | I | V | V | V | L | TMD3 | Conservative |
| **Site** | G1 | G1A | G1B | G2 | G3 | G4 | Out | **Location** | **Substitution type** |
| **Non-structural protein 2B (continued)** | | | | | | | | | |
| 116 | L | L | V | L | L | L | I | TMD3 | Conservative |
| **Non-structural protein 3** | | | | | | | | | |
| 11 | K | K | K | R | K | K | K | Protease | Conservative |
| 33 | R | R | R | S | R | R | R | Protease | Non-conservative |
| 71 | N | N | N | S | N | N | N | Protease | Conservative |
| 106 | A | A | A | V | A | A | A | Protease | Conservative |
| 147 | I | I | I | V | I | I | V | Protease | Conservative |
| 355 | Y | Y | C | Y | Y | Y | Y | NTP-ase helicase | Conservative |
| 368 | M | M | M | I | M | M | M | NTP-ase helicase | Conservative |
| 445 | S | S | S | G | S | S | G | NTP-ase helicase | Conservative |
| 515 | D | D | D | E | D | D | D | NTP-ase helicase | Conservative |
| 562 | K | K | K | R | K | K | K | NTP-ase helicase | Conservative |
| 583 | I | I | I | T | I | I | I | NTP-ase helicase | Non-conservative |
| 590 | K | K | K | R | K | K | K | NTP-ase helicase | Conservative |
| **Non-structural protein 4A** | | | | | | | | | |
| 17 | A | A | A | V | A | A | A | A1 | Conservative |
| 58 | A | A | A | A | V | A | A | A4 TM1 | Conservative |
| 62 | V | V | V | I | V | V | V | A4 TM1 | Conservative |

| | | | | | | | | | |
|---|---|---|---|---|---|---|---|---|---|
| 71 | M | M | M | L | M | M | M | A4 TM1 | Conservative |
| 88 | V | V | V | A | V | V | C | A5 TM2 | Conservative |
| 90 | A | A | A | V | A | A | A | A5 TM2 | Conservative |
| **Non-structural protein 4B** | | | | | | | | | |
| 11 | T | T | T | D | T | T | E | A1 | Non-conservative |
| 14 | R | R | R | K | R | R | R | Loop between a1 and a2 | Conservative |
| 25 | N | N | N | T | N | N | N | Loop between a1 and a2 | Conservative |
| 30 | P | P | P | S | P | P | M | Loop between a1 and a2 | Non-conservative |
| 68 | V | V | I | V | V | V | V | A3 pTMD2 | Conservative |
| 118 | T | T | T | A | T | T | V | A5 pTMD3 | Non-conservative |
| **Site** | G1 | G1A | G1B | G2 | G3 | G4 | Out | **Location** | **Substitution type** |
| **Non-structural protein 4B (continued)** | | | | | | | | | |
| 188 | A | A | A | S | A | A | A | A7 pTMD4 | Non-conservative |
| 189 | V | V | V | L | V | V | L | A7 pTMD4 | Conservative |
| 195 | I | I | I | V | I | I | V | A8 pTMD4 | Conservative |
| 214 | D | D | D | E | D | D | D | A8 | Conservative |
| 227 | T | T | I | T | T | T | T | A9 pTMD5 | Non-conservative |
| 252 | E | E | E | E | E | G | E | A9’ pTMD5 | Non-conservative |
| 256 | F | F | F | F | F | C | F | Loop after a9’ | Non-conservative |
| **Non-structural protein 5** | | | | | | | | | |
| 26 | S | S | S | N | S | S | Q | Capping 2 OMTase | Conservative |
| 91 | A | A | A | T | A | A | A | Capping 2 OMTase | Non-conservative |
| 99 | E | E | D | E | E | E | E | Capping 2 OMTase | Conservative |
| 156 | I | I | I | V | I | I | V | Capping 2 OMTase | Conservative |
| 168 | V | V | V | V | V | T | V | Capping 2 OMTase | Non-conservative |

| | | | | | | | | | |
|---|---|---|---|---|---|---|---|---|---|
| 194 | K | K | K | R | K | K | K | Capping 2 OMTase | Conservative |
| 274 | T | T | T | S | T | T | S | Loop between Capping 2 OMTase and RdRp | Conservative |
| 277 | K | K | K | R | K | K | K | Loop between Capping 2 OMTase and RdRp | Conservative |
| 283 | I | I | I | V | I | I | V | Loop between Capping 2 OMTase and RdRp | Conservative |
| 512 | A | A | A | A | A | S | A | RdRp RNA binding site | Non-conservative |
| 587 | A | A | A | T | A | A | A | RdRp | Non-conservative |
| 640 | I | I | I | V | I | I | L | RdRp | Conservative |
| 665 | V | V | I | V | V | V | V | RdRp conserved polymerase motif C | Conservative |
| 721 | V | V | V | L | V | V | L | RdRp | Conservative |
| 836 | S | S | S | A | S | S | T | RdRp | No-conservative |
| **Site** | G1 | G1A | G1B | G2 | G3 | G4 | Out | **Location** | **Substitution type** |
| **Non-structural protein 5 (continued)** | | | | | | | | | |
| 873 | N | N | N | Y | N | N | N | RdRp | Conservative |
| 878 | V | V | V | I | V | V | I | RdRp | Conservative |
| 889 | Q | Q | Q | L | Q | Q | Q | Loop after RdRp | Non-conservative |
| 898 | H | H | H | H | N | H | T | Loop after RdRp | Non-conservative |

#### Supplementary Figure 1

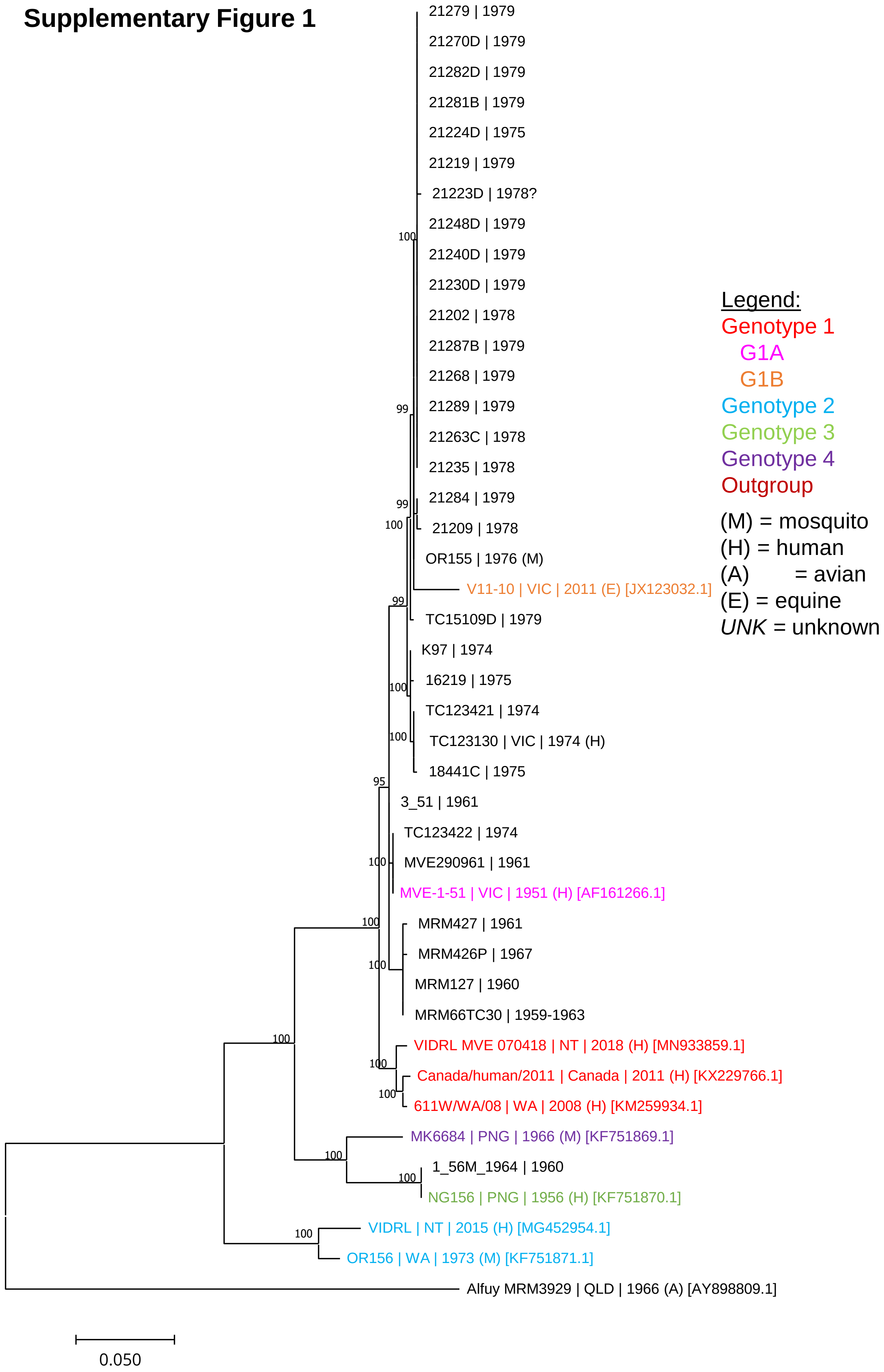

### Supplementary Figure 2

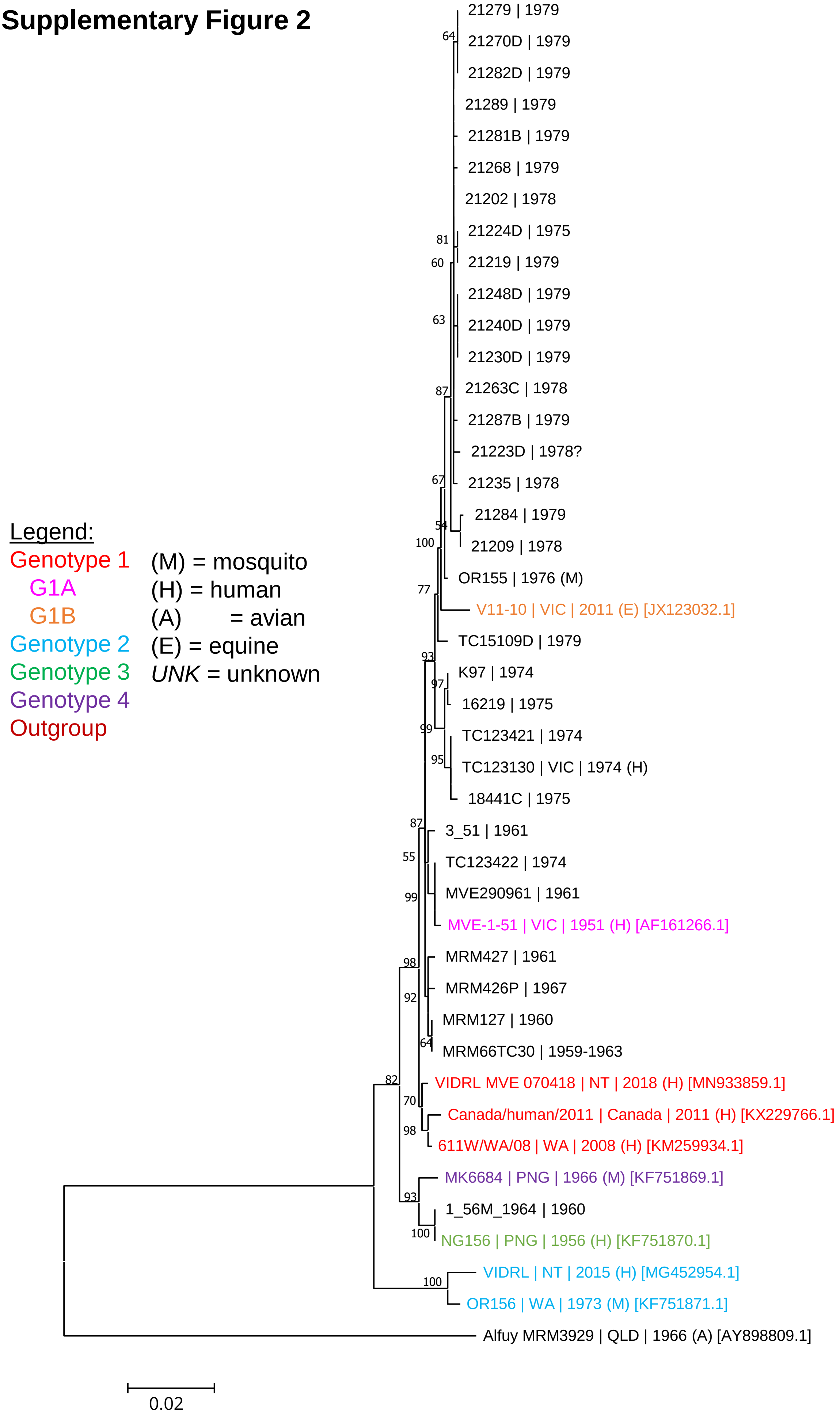
